# Single-cell dissection of cervix and placenta reveal both novel and overlapping cell types

**DOI:** 10.1101/2025.07.09.663838

**Authors:** Margot van Riel, Lore Lannoo, Anne Pexsters, Olga Tsuiko, Ilse Parijs, Irene Claes, Thierry Voet, Dirk Timmerman, Joris R. Vermeesch

## Abstract

Placental trophoblasts have been detected in cervical smears early in gestation, creating opportunities for non-invasive prenatal diagnosis. However, trophoblast isolation is limited by lack of a cell catalogue and molecular profile of cervical smears. To establish an atlas and explore the potential of single-cell RNA-sequencing to detect cervical trophoblasts, 10,539 single-cell transcriptomes from 12 non-invasive exocervical smears from pregnant women and 34,565 cells from six placentas were profiled. We uncovered a novel extravillous trophoblast cell subtype characterized by epithelial marker genes and reduced HLA-G expression. Integration of both cell atlases demonstrated surprisingly similar expression profiles between maternal epithelial cells and placental extravillous trophoblasts, indicating trophoblasts retained epithelial properties without an invasive mesenchymal phenotype. Differential expression analysis identified novel markers discriminating cervical and placental cell types. Using those markers for immunocytochemistry we demonstrate the frequency of exocervical trophoblast cells to be lower than reported.

## Introduction

The cervix is a complex, heterogeneous organ which undergoes extensive changes throughout pregnancy^1^. These include immature squamous metaplasia of the epithelium and hyperplasia of the endocervical glands, which coincides with an increase in cervical mucus^2,3^. In addition to the epithelial transformations, the cervical immune environment is altered during pregnancy, and plays a key role in the protection against infections^4,5^. Cervical smears provide an elegant way to investigate the cervix and are generally applied to detect (pre-)cancerous lesions^6^. Interestingly, placental trophoblast cells have been observed in cervical mucus of pregnant women, which could lead to an alternative source of placental cells for prenatal diagnosis^7^.

Trophoblast cells cover the placental villi, which form the functional unit of the placenta. A well-developed and functioning placenta is indispensable for a successful pregnancy as it performs similar functions to the lungs, liver, gut, kidneys and the endocrine glands for the developing fetus^8^. The placental villous is characterized by a mesenchymal core, surrounded by villous cytotrophoblast cells (VCTs). VCTs are a precursor cell population that gives rise to more specialized trophoblast subtypes: the syncytiotrophoblast (SCT) and extravillous trophoblasts (EVTs)^9^. The SCT is a multinucleated layer that covers the entire placental villous and is the main site of exchange for nutrients and waste products between mother and fetus. Furthermore, the SCT is the major site of placental hormone production^9^. There are two types of placental villi; floating villi float freely in the intervillous space, while anchoring villi are attached to the uterus. At the tips of these anchoring villi, trophoblasts proliferate in cell columns and EVTs arise. EVTs are invasive cells that populate different structures of the decidua, including the interstitium, the spiral arteries, the uterine veins and lymphatic vessels, and the decidual glands^10^.

Recent studies investigating cervical trophoblasts report the presence of both SCTs and EVTs^11,12^. To date, there are two hypothesized routes that invasive EVTs might take to reach the cervical environment. First, they could replace the uterine epithelium at the border of the placenta. Similarly, ulcerations of the decidua might permit release of syncytial fragments^13,14^. Second, by invasion of the uterine glands, EVTs could be expelled with glandular secretions into the uterus and further towards the cervix^13^. From the cervix, these cells could be obtained non-invasively, avoiding invasive procedures such as chorionic villous sampling or amniocentesis^15^. Current studies mainly target EVTs for isolation, which represent only a subset of all cervical trophoblast cells^11,16,17^.

The composition of cervical smears has been described based on cytological examination and flow cytometry analysis. The former focuses on epithelial cells for cancer detection, and the latter explores the immune cells residing at the female reproductive tract^3,18,19^. Over recent years, single-cell RNA-sequencing (scRNA-seq) has allowed for the disentanglement of the cellular compositions and molecular characteristics of complex tissues, providing a more fine-grained analysis as compared to cytology. For example, placental cell atlases have identified different subpopulations of trophoblast cells^20,21^. Here, we characterize the cellular composition of the exocervical environment and explore the presence of cervical trophoblasts using large-scale single-cell RNA-sequencing. We provide a molecular map of exocervical smears and early gestational placentas and identify distinct cell populations and subtypes. Moreover, we identify a novel extravillous trophoblast subtype, characterized by epithelial signatures. Integration of both cell atlases showed similarities between placental trophoblasts and cervical epithelial cells that pinpoint an extra challenge for cervical trophoblast identification and isolation.

## Results

### Single-cell atlas of exocervical smears from pregnant women

Twelve exocervical smears were collected. After quality control and sample integration, scRNA-seq profiles from 10,973 cells were retained, with a cell count per sample ranging from 53 to 1,813 cells (Supplementary Table 1). Using graph-based clustering, 16 different clusters were identified. As both immune and epithelial cell types were expected, these were first distinguished by expression of *PTPRC* (encoding CD45) and *KRT13*, respectively (Figure 1a). Clusters 9, 10 and 14 were defined by pronounced expression of ribosomal, mitochondrial, and heat shock genes, respectively, indicative of apoptotic cells or cells under stress. These clusters were removed, leaving 10,539 cells for downstream analyses (Figure 1b). Differences in the proportions of cell types were noted between samples (Figure 1c, Supplementary Table 2).

**Fig. 1.**
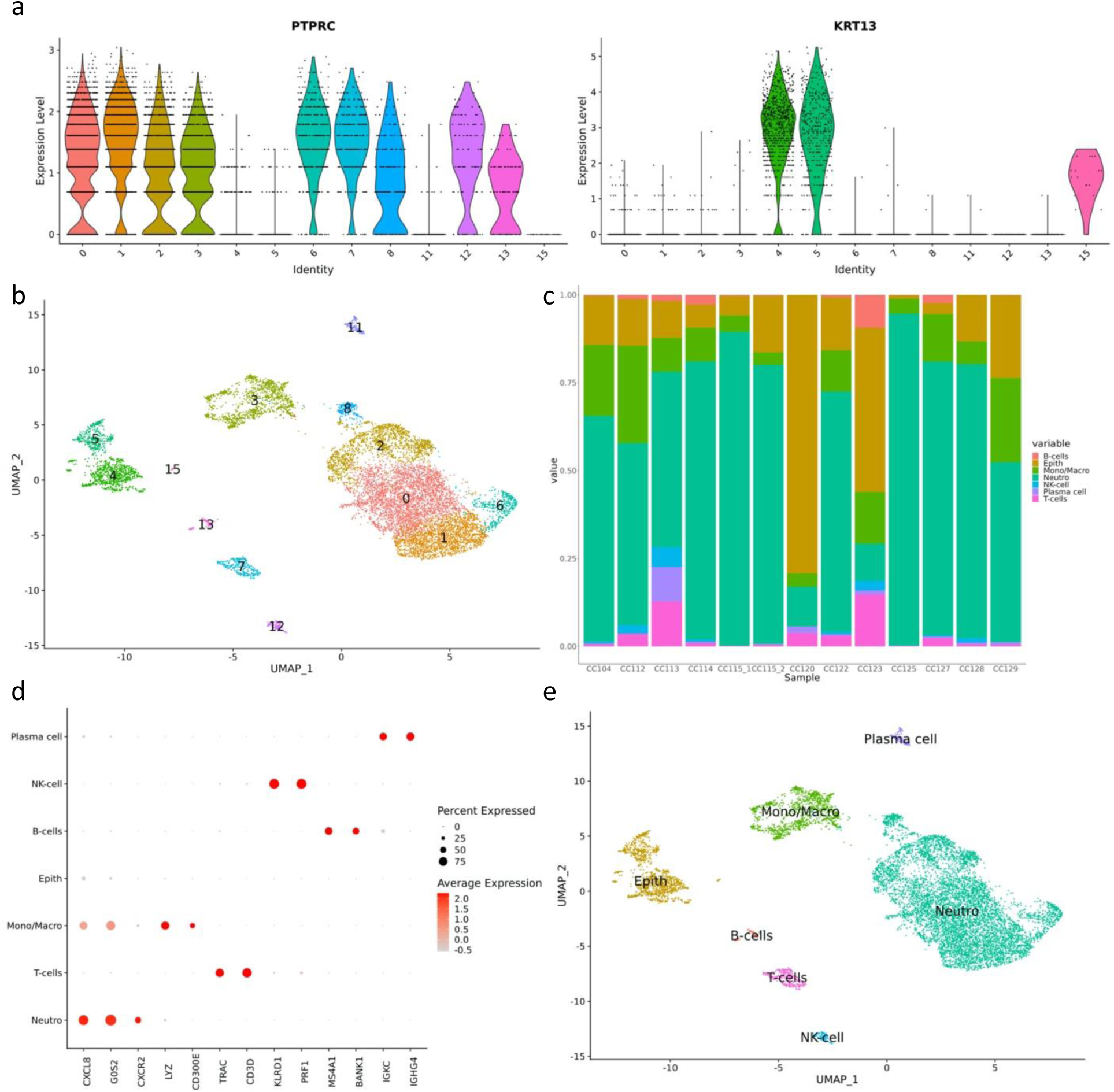
Single-cell analysis of exocervical samples during early pregnancy. a. Violin plots with expression of *PTPRC* and *KRT13* per cluster. b. UMAP after graph-based clustering at resolution 0.3. c. Bar plot showing the contribution of each sample to each cluster after sample integration. d. Dot plot showing immune cell sublineage markers. e. UMAP with annotated cell types. Epith = epithelial cells, Mono/Macro = monocytes and macrophages, Neutro = Neutrophils.

Based on marker gene analysis, we could identify a large proportion of neutrophils, monocytes and macrophages, T-cells, plasma cells, Natural Killer-cells, and B-cells (Figure 1d, e). Neutrophils were further subdivided into five corresponding clusters, based on distinct gene expression (Figure 2a, Table 1). Cluster 13, identified as B-cells, was spread over two groups of cells (Figure 1b). A subset of this cluster was found to consist of plasmacytoid dendritic cells, characterized by *LILRA4*, *IRF7*, *PACSIN1* and *TCF4* expression with no expression of the B-cell specific markers *MS4A1* and *BANK1* (Supplementary Fig. 1).

**Fig. 2.**
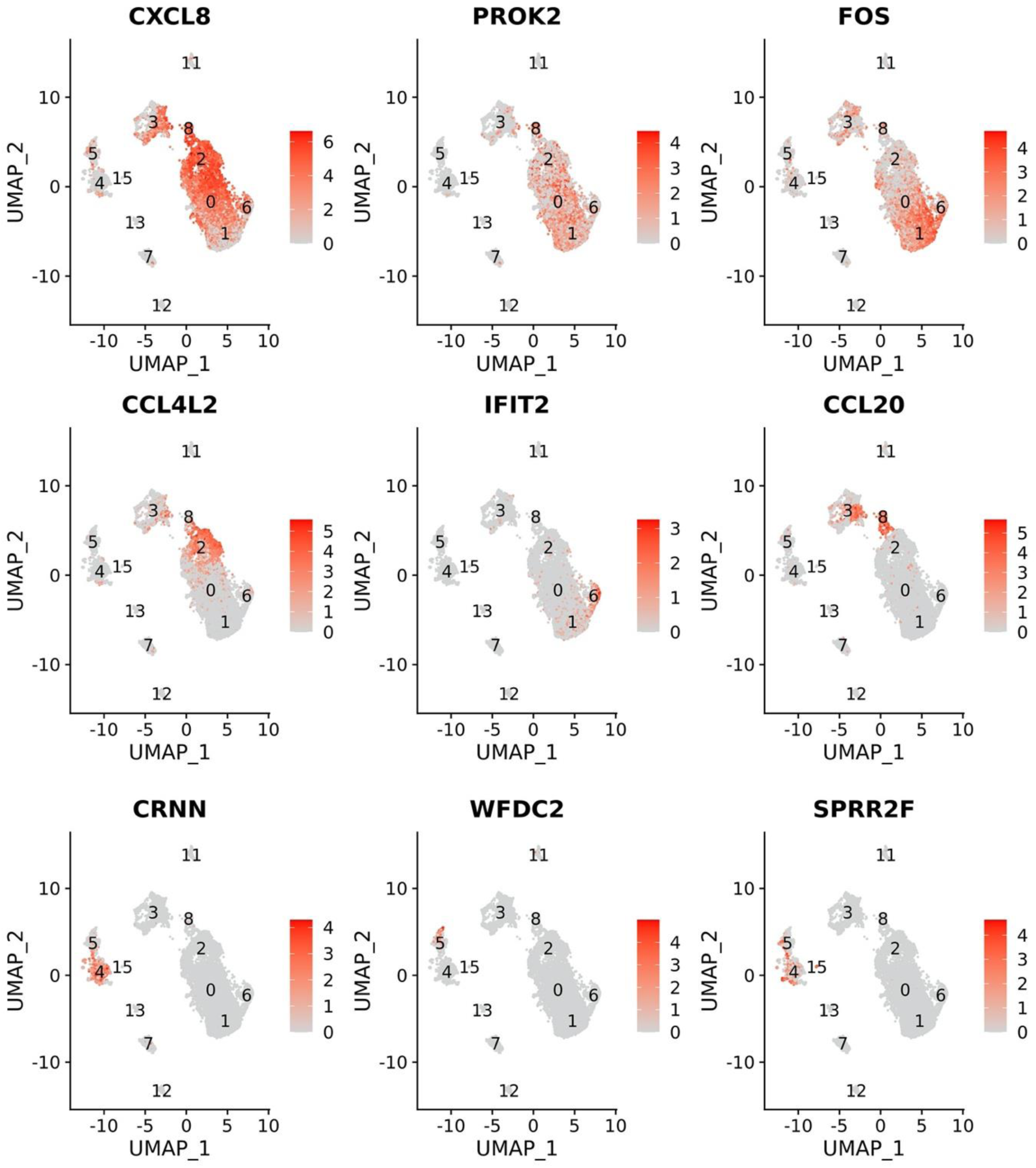
Feature plots showing expression of marker genes of different exocervical cell populations. . (a) CXCL8 is a general neutrophil marker, the other five plots depict expression of the top expressed gene of each neutrophil cluster, in accordance with Table 1. (b) Epithelial cell clusters 4 and 5 are characterized by *CRNN* and *WFDC2*, markers of squamous and glandular epithelial cells, respectively, while cluster 15 is marked by expression of *SPRR* genes.

**Table 1:**
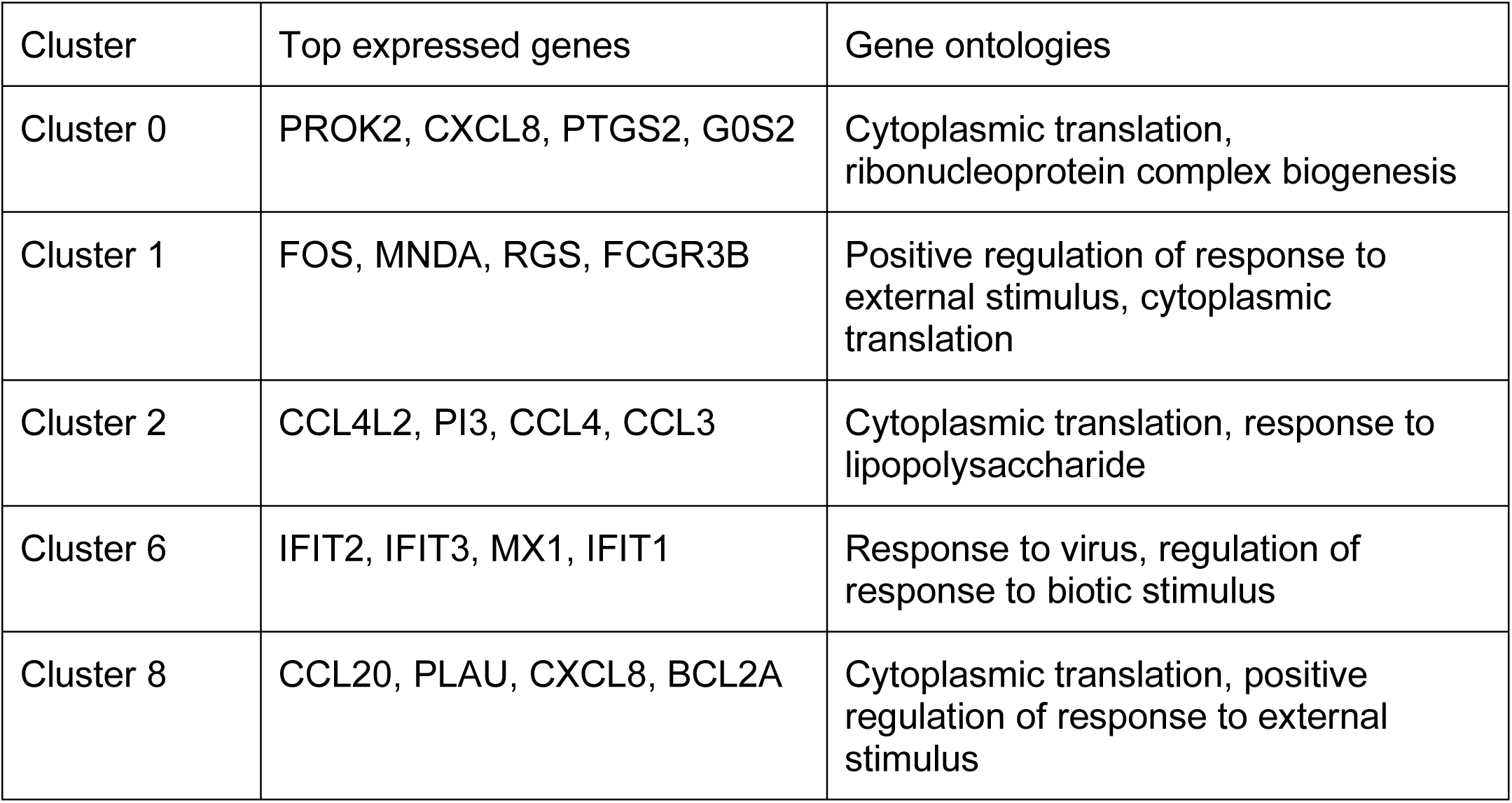
Subpopulations of neutrophils in non-invasive exocervical samples.

Three clusters were of epithelial origin, and characterized by expression of genes of the epidermal differentiation complex (EDC) (Figure 1a, Figure 2b). The EDC regulates the terminal differentiation program of keratinocytes, which is important for epithelial tissue development and repair. A cornified envelope is formed at the end of the differentiation trajectory, consisting of mitotically inactive flattened cornified cells^22^. The largest epithelial population (cluster 4) highly expresses EDC genes, such as *CRCT1*, *LCE3D, SPRR3* and *CNFN*, which are expressed in the squamous epithelia^23–26^. Another subset of epithelial cells (cluster 5) shares expression of EDC genes, but also shows upregulation of glandular epithelial cell-specific genes: *S100P*, *WFDC2* and *MUC4*. A small cluster (cluster 15) is derived from a single sample and all cells are predicted to be in the G2M phase of the cell cycle, suggesting cell proliferation. The gene expression profile of these cells is similar to the squamous cells in cluster 4, but with higher expression of the *SPRR* genes that are part of the cornified envelope precursor family of the EDC complex, as well as genes involved in immune response and mucosal defense. Additionally, there was lower expression of *CRNN*, which plays a role in epithelial homeostasis, and of *PSCA,* which might have inhibitive cell proliferation capacities^27^. Therefore, these cells might represent a less differentiated state of cervical squamous epithelial cells.

### Single-cell atlas of early placental material

Six placental samples with gestational ages ranging from 9.7 to 12.6 weeks were collected by hysteron-embryoscopy. Five placenta samples were aneuploid and one was euploid (Supplementary Table 3). A total of 34,941 cells passed quality control, with a cell count per sample ranging from 1,431 to 10,697. Different cell types were identified based on expression of cell-specific marker genes (Figure 3a, b; Supplementary Table 4)^20,21,28^. In total, graph-based clustering identified 19 clusters (Figure 3c), however, cluster 15 lacked clear marker genes, as upregulated genes were long non-coding RNAs and RNA genes. Therefore, this cluster was removed from further analysis, leaving a final dataset of 34,565 placental cells (Figure 3d, Table 2). Fibroblasts were the most prevalent cell type, comprising 48% of all cells, followed by macrophages (19%), villous cytotrophoblasts (VCTs) (17%), extravillous trophoblasts (EVTs) (12%), syncytiotrophoblasts (SCTs) (4%), and vascular endothelial cells (VECs) (1%). Two populations of macrophages were detected, and genotype analysis using freemuxlet demonstrated one was fetal in origin (Hofbauer cells; constituting 15% of all cells) and the other maternal (4%) (Figure 3e).

**Fig. 3.**
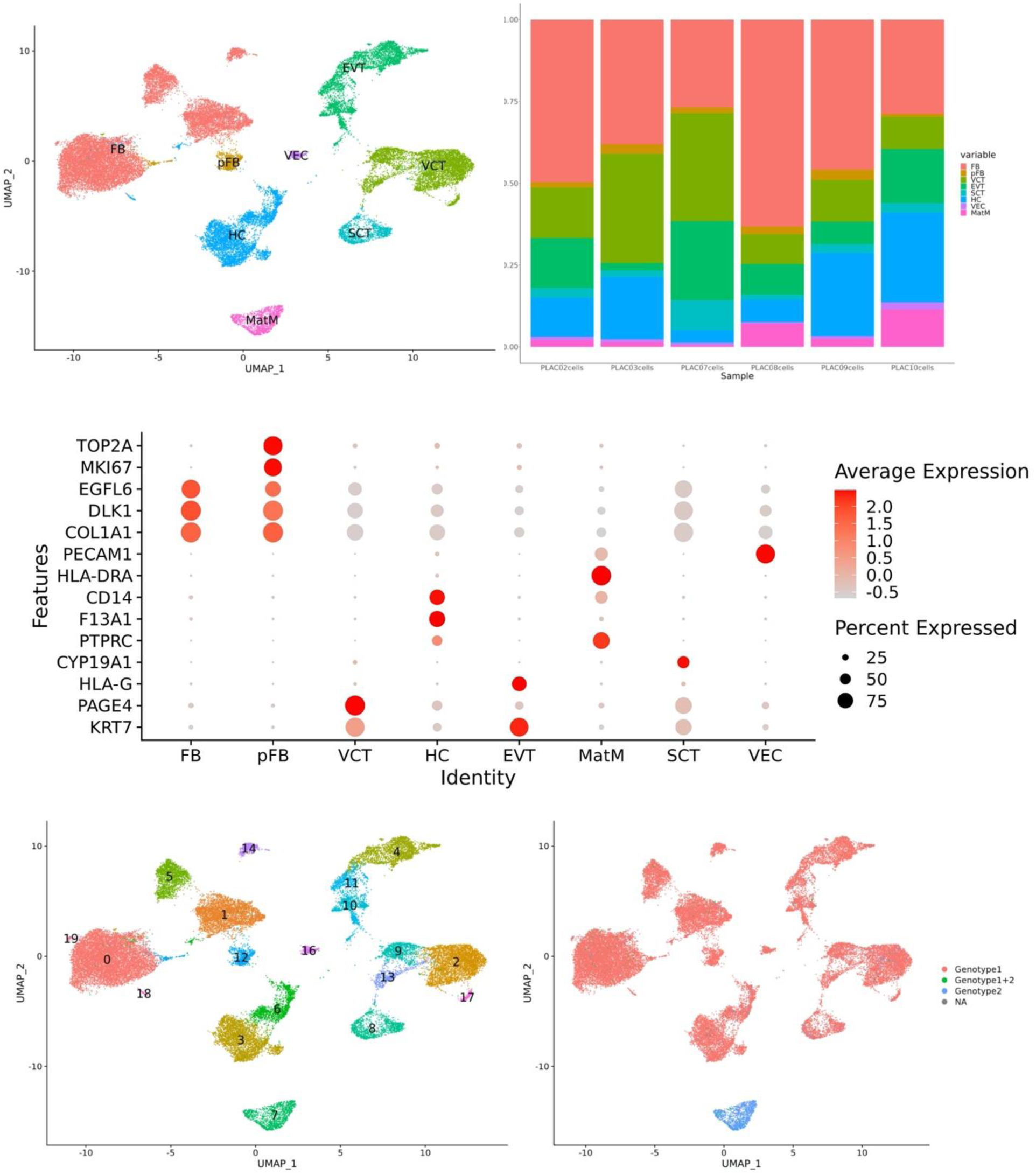
Single-cell analysis of the early placenta. **a.** UMAP with cell type annotations based on established marker genes. EVT = extravillous trophoblasts, FB = fibroblasts, HC = Hofbauer cells, MatM = maternal macrophages, pFb = proliferating fibroblasts, SCT = syncytiotrophoblast, VCT = villous cytotrophoblast, VEC = vascular endothelial cells. b. Dot plot showing expression of established marker genes per cell type. c. Bar plot showing the contribution of each sample to each cluster after sample integration. d. UMAP after graph-based clustering at resolution 0.3, yielding 19 clusters. e. UMAP with the BEST.GUESS results of the freemuxlet analysis, which indicated a minor fraction of cells having a different genotype in each sample, clustering together. Genotype 1 represents a first cell population, Genotype 2 a second and Genotype 1+2 indicates a doublet of Genotype 1 and 2. NA means freemuxlet gave no result.

**Table 2:** Subpopulations of placental cells.

| Annotation | Cluster | Top expressed genes | Gene ontologies |
| --- | --- | --- | --- |
| Trophoblast cells |  |  |  |
| VCT | 2 | PAGE4, PEG10, PHLDA2, XAGE3 | Generation of precursor metabolites and energy, energy derivation by oxidation of organic compounds |
| VCT | 9 | PEG10, PAGE4, IFI6, EGFR | Generation of precursor metabolites and energy, proteasomal protein catabolic process |
| VCT | 13 | CGA, KRT23, HSD17B1, CYP19A1 | Proteasomal protein catabolic process, generation of precursor metabolites and energy |
| VCT | 17 | PAGE4, PHLDA2, XAGE3, IFI6 | Generation of precursor metabolites and energy, energy derivation by oxidation of organic compounds |
| EVT | 4 | AOC1, NOTUM, HPGD, PAPPA2 | Actin filament organization, ameboidal-type cell migration |
| EVT | 10 | GABRP, KRT17, TACSTD2, CADM1 | Regulation of actin filament-based process, ameboidal cell-type migration |
| EVT | 11 | KRT19, CCNE1, FABP7, KRT7 | Organelle fission, mitotic cell cycle phase transition |
| SCT | 8 | CGA, KISS1, CSH1, TFPI2 | Multi-multicellular organism process; reproductive structure development |
| Villous core cells |  |  |  |
| FB | 0 | CXCL14, CFD, PTGDS, PLPP3 | Ossification, mesenchyme development |
| FB | 1 | POSTN, COL1A2, COLA1A, COL3A1 | Extracellular matrix organization, extracellular structure organization |
| FB | 5 | TAGLN, ACTG2, ACTA2, DES | Wound healing, extracellular matrix organization |
| FB | 12 | TOP2A, CENPF, HMGB2, UB2C | Organelle fission, nuclear division |
| FB | 14 | PLA2G2A, AREG, REN, COX4I2 | Cell-substrate adhesion, wound healing |
| FB | 18 | TAGLN, ACTA2, TPM2, CRYAB | Aerobic respiration, cellular respiration |
| FB | 19 | VIM, IGFBP3, IFITM3, LUM | Viral process, modulation by symbiont of entry into host |
| HC | 3 | SPP1, F13A1, RNASE1, CD14 | Immune-response regulating pathway, actin filament organization |
| HC | 6 | TMSB4X, AIF1, F13A1, DAB2 | ATP metabolic process, generation of precursor metabolites and energy |
| MatM | 7 | HLA-DRA, APOC1, LYZ, CD74 | T-cell activation, positive regulation of cytokine production |
| VEC | 16 | PECAM1, EGFL7, CLDN5, KDR | Ameboidal-type cell migration, cell junction assembly |

As a control for our dataset and methodologies, we integrated our atlas with another published placental atlas^28^. Similar cell populations were detected (Supplementary Data), highlighting that aneuploidies do not detectably affect the overall transcriptome clustering.

Four VCT clusters (2, 9, 13 and 17) were identified (Table 2). Cluster 17 was derived entirely from sample PLAC03. While cluster 2 was characterized by general VCT markers only, cluster 9 showed additional upregulation of genes involved in organization of the extracellular matrix, such as *EGFR*, *ITIH5*, *SPINT1* and *FBN2*, compared to other VCT clusters. *EGFR* has been associated with cell cycle progression in VCTs^29,30^. Part of the cells in cluster 9 are indeed proliferating, as confirmed by expression of cell proliferation markers *MKI67* and *TOP2A* (Supplementary Fig. 2). Cluster 13 is characterized by expression of genes involved in hormone production. Moreover, these cells express *ERVFRD-1,* which encodes for Syncytin-2, a protein that was found to induce cell fusion of human cells^21^. Gene ontology analysis showed that this cluster also plays a role in macro-autophagy. Autophagy activity at the placenta was previously shown to take place in the SCT^31^. There are three EVT clusters (4, 10, 11) (Figure 4). Marker genes of cluster 4 are associated with epithelial-mesenchymal transition (EMT) and cellular invasion^28,30^.

**Fig. 4.**
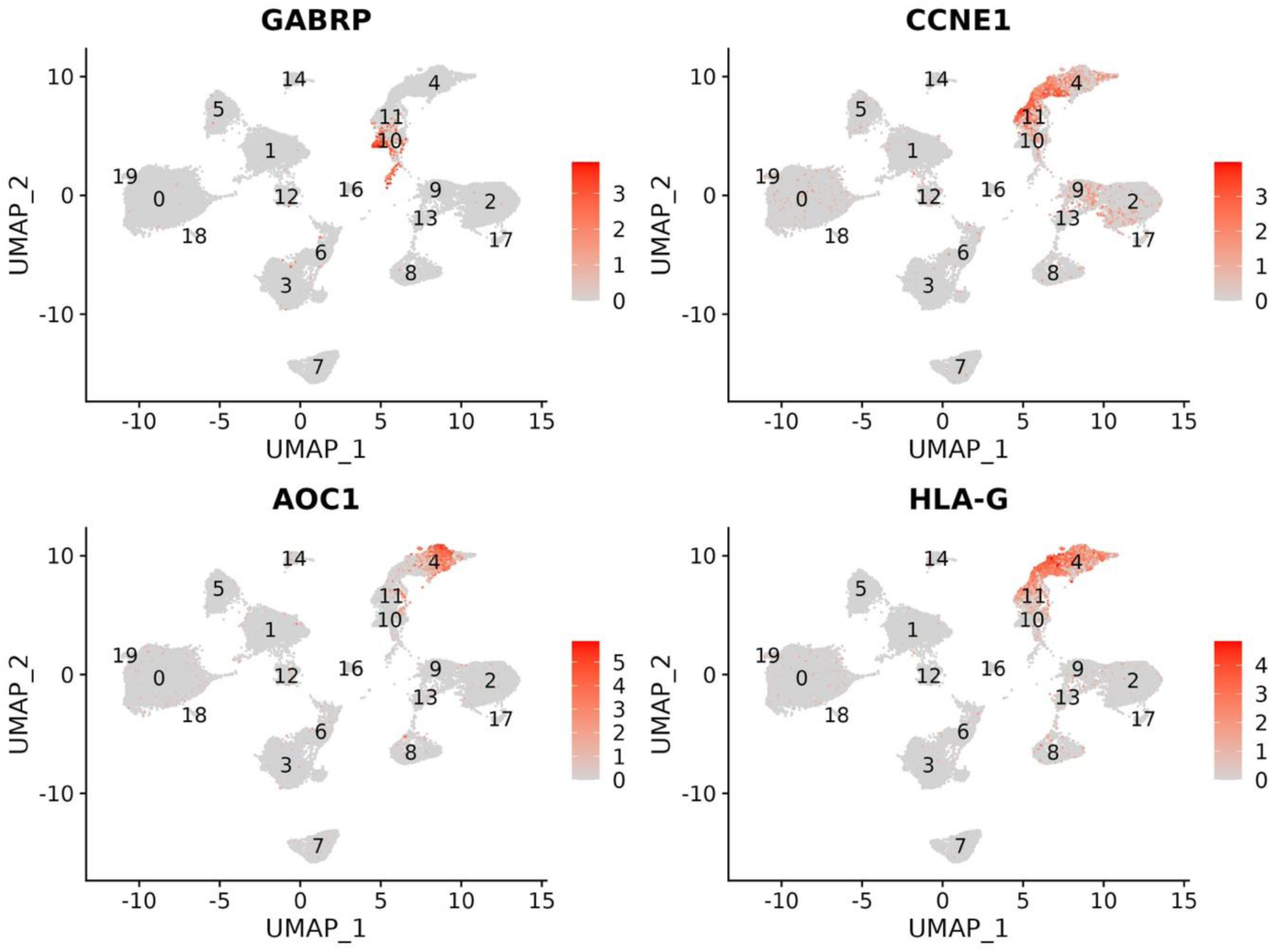
Feature plots showing markers of the different EVT subtypes detected in placental samples. Cluster 10 has marked expression of epithelial marker *GABRP*, while cluster 11 is characterized by strong proliferation, as indicated by cyclin gene *CCNE1*. *AOC1* is a marker gene of cluster 4, which also has the highest expression of *HLA-G*.

Furthermore, there is the highest expression of markers of terminally differentiated EVTs, such as *HLA-G* and *AOC1*, which was shown to be expressed by EVTs in close proximity to decidual vessels^32^. Cluster 10 has the lowest percentage of HLA-G expressing cells of all three EVT clusters, and showed high expression of genes pointing to an epithelial cell state, such as the cytokeratin genes, *GABRP* and *TACSTD2*. Freemuxlet analysis indicated the genotype of these cells to be the same as other placental populations, ruling out contamination by decidua-derived cells (Figure 3e). Cluster 11 expresses cytokeratin genes as well, but also genes related to more differentiated EVTs such as *LAIR2*, associated with spiral artery remodeling, and *DIO2*, a regulator of trophoblast invasion^33,34^. Additionally, this cluster was marked by cell proliferation as shown by expression of cyclin genes. On the other end of the trophoblast differentiation trajectory are the SCTs, which are grouped in cluster 8.

To reconstruct the paths that the trophoblast undergo through different states, trajectory analysis was performed. These results show three distinct trajectories, starting from the VCT progenitor cells, to (1) the invasive EVTs, (2) the SCTs and (3) the EVTs with an epithelial phenotype (Supplementary Figure 3).

The fibroblasts residing in the mesenchymal core of the placental villous are the largest population of cells with seven distinct clusters (Table 2, Supplementary Fig. 4). One (cluster 0) is characterized by expression of pro-inflammatory genes and decreased expression of *THY1*, which is associated with increased differentiation of mesenchymal stromal cells^35^. Two other clusters are characterized by the expression of extracellular matrix proteins (cluster 1) and muscle proteins (cluster 5). As mentioned, there is a subpopulation of proliferating fibroblasts (cluster 12). Marker genes of the last main cluster (cluster 14) are implicated in phospholipid metabolism and regulation of blood pressure, sodium and fluid homeostasis. Two smaller clusters (cluster 18 and cluster 19) were largely derived from single samples; it remains unclear whether their existence is related to the placental aneuploidy of their specific sample source or whether they contain cell doublets with other cell types.

Two Hofbauer cell clusters were observed. Both contained proliferating cells (Supplementary Fig. 2). The larger one (cluster 3) expressed *CCL3*, *CCL4* and *CXCL8*, which are proangiogenic factors, consistent with their role in placental vasculogenesis and angiogenesis^36^. Cluster 6 showed reduced expression of these proangiogenic genes but lacked specific marker genes (Supplementary Fig. 5). Maternal macrophages (cluster 7) share expression of *SPP1* with the Hofbauer cells, but *F13A1* and *LYVE1* expression is unique to Hofbauer cells (Supplementary Fig. 6). The small population of VECs were characterized by high expression of endothelial-specific genes.

### No trophoblast cells were detected in exocervical smears based on scRNA-seq analyses

We performed several analyses to explore the presence of trophoblast cells in the exocervical smears. First, marker genes of all clusters were investigated. There was no cluster characterized by trophoblast marker gene expression. Second, expression of trophoblast-specific genes was explored. There were no cells expressing *HLA-G* and *CYP19A1*, markers specific of EVTs and SCTs, respectively (results not shown). Third, expression of the Y-chromosomal *RPS4Y1* gene was checked, as six pregnant women carried a male fetus (Supplementary Table 1). While expression of the Y-chromosomal gene *RPS4Y1* is observed in all fetal clusters of the placental cell atlas, its expression was not detected in any of the exocervical cells. Lastly, the genotype of all cells was analyzed, as this differs between placental trophoblast cells and maternal cells. Mixtures of genotypes can be disentangled from expression data using freemuxlet, which identified a second genotype in ten of the 11 exocervical smears. Overall, 0%-25.3% of cells per sample had a different genotype (Supplementary Table 5). In total, this corresponds to 555 cells out of 9,112 cells (6.1%) with a freemuxlet result, which is much higher than the reported frequency of 0.05% of placental cells in a cervical smear^37^. An alternative interpretation could be that these are sperm cells. However, previously identified sperm cell markers (*PRM1*, *PRM2*, *TSSK6*) were not expressed in any of the cells^38^. Moreover, cells with a different genotype were scattered over the different clusters instead of clustering together (Supplementary Fig. 7). In conclusion, there is no indication of the presence of placental trophoblast cells in our dataset of exocervical smears.

### Placental EVTs resemble exocervical epithelial cells

To explore similarities and differences amongst placental and exocervical cells, we integrated our placental and exocervical cell atlases (Figure 5). As expected, most clusters consist of either placental or exocervical cells. However, there are some notable exceptions. Unexpectedly, a subset of the exocervical epithelial cells co-cluster with placental cells. This co-clustering consists of exocervical epithelial cells and EVTs: the novel EVT subtype we identified (cluster 10) makes up 67% of all placental cells co-clustering with the epithelial cells (Figure 5c). The intermediate EVT subtype (cluster 11) is responsible for 29%, and the remaining 4% is divided over the different other clusters. Conserved markers between the placental and cervical cells co-clustering were explored (Supplementary Fig. 8). This integrated analysis demonstrates the resemblance of placental cells with the epithelial cells of the cervical canal. Co-clustering additionally occurs between the placental immune cells and cervical macrophages, and to a lesser extent Natural Killer- and T-cells, and neutrophils. Cervical immune cells also co-cluster with the Hofbauer cells, possibly due to shared expression of macrophage markers.

**Fig. 5.**
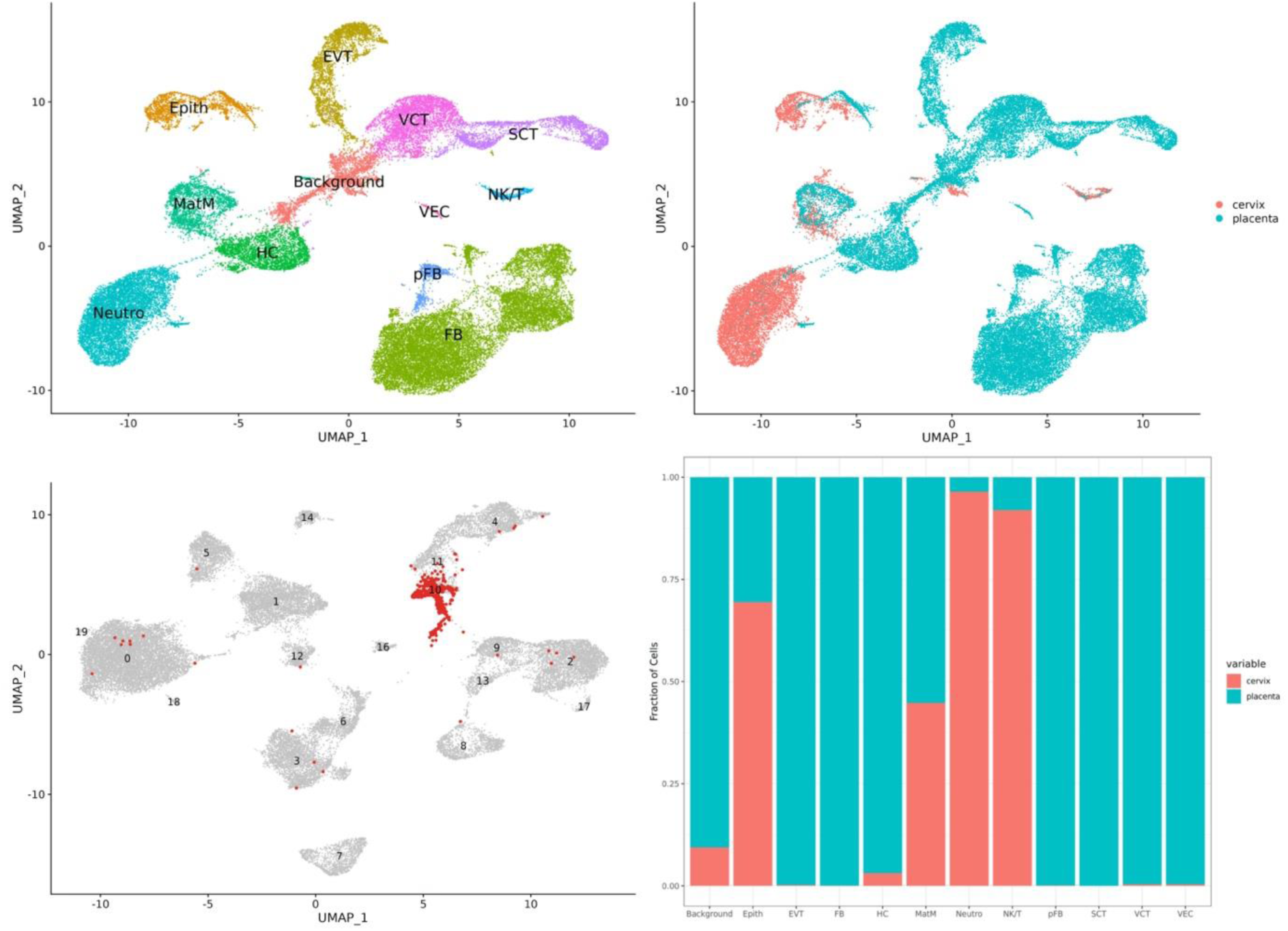
Integrated analysis of the placental and exocervical datasets. **a.** UMAP of the integrated placental and exocervical dataset with cell annotations based on marker genes. The cluster labeled as ‘Background’ lacked specific marker genes; their cell annotations from the individual clustering results are noted in Supplementary Table 6. b. UMAP of the integrated placental and exocervical dataset with sample origin plotted, indicating the presence of clusters derived from cervix or placenta only, as well as mixed clusters. c. UMAP of the placental dataset highlighting the cells that co-cluster with the epithelial cells. d. Bar plot showing the relative contribution of each sample type to each annotated cell type.

### Identification of markers for cell selection

The isolation of cervical trophoblasts mainly relies on *HLA-G*, which is only expressed by a subset of trophoblasts and is insufficient to obtain a pure trophoblast population^39,40^. As our analyses did not detect trophoblast cells in the exocervical dataset, we explored differentially expressed genes between maternal and trophoblast cells to identify new markers. We searched for genes expressed by (1) EVTs only, demonstrated to reside in the cervix and the main target^12^, (2) SCTs only, as these cells have been reported in exocervical smears as well^11^, and (3) epithelial cells only, as potential targets for negative selection. Only genes expressed on the cell membrane were selected. Seven markers unique for EVTs and absent in exocervical cell populations were detected, however, others were found to be expressed in epithelial or immune cell populations as well (Supplementary Table 7). Three additional membrane markers were expressed by SCTs but absent in the exocervical cell populations (Supplementary Table 8), and 13 markers were uniquely expressed by the epithelial cells and absent in the EVT and SCT populations (Supplementary Table 9). Hence, besides *HLA-G*, *ADAM12*, *MCAM*, and *PSG2* present as potential targets for positive selection of trophoblast cells while *MAL*, *RHCG* and *MUC21*, targeting epithelial cells, could be applied for negative selection.

### Immunostaining shows the presence of cervical trophoblasts, at frequency lower than previously reported

We first tested the antibodies targeting trophoblast cells on chorionic villous samples (CVS). Four trophoblast-specific antibodies were explored (Supplementary Table 10): *HLA-G*, *ADAM12*, *MCAM* and *PSG2*. The genes targeted by these antibodies are all expressed by EVTs, and *PSG2* is additionally expressed by SCTs^41^. The frequency of EVTs was first evaluated by counting the fraction of *HLA-G* positive cells in a CVS, which was 13.4%. Then, a single CVS was divided over three parts in which *HLA-G* was multiplexed with *ADAM12*, *MCAM* and *PSG2*. Combined *HLA-G* and *ADAM12*, *MCAM* or *PSG2* signals were identified in 21.4%, 18.6% and 27% of cells, respectively, confirming specificity of these antibodies for a subset of placental cells. In addition, out of the 162 counted cells in the sample part labeled with *MCAM* and *HLA-G*, one cell was identified that was only positive for *HLA-G* and one cell was only positive for *MCAM*. The combinations of *HLA-G* and *ADAM12* or *PSG2*, only revealed double-labeled cells (Supplementary Fig. 9).

In parallel with cervical samples from pregnant women, three non-pregnant control samples were collected and stained with *HLA-G* combined with one of the three different potential trophoblast-specific markers. In the sample labeled with *HLA-G* and *ADAM12* combined, a single *ADAM12*-positive - but *HLA-G* negative - cell was identified. In the other samples, no positive fluorescent cells were identified.

In cervical samples of pregnant women, the fraction of trophoblast cells was previously reported to be 1 in 2,000^37^. Again, the antibodies targeting trophoblast cells were combined with the *HLA-G* antibody. Approximately one fifth of the final stained cell suspension was analyzed. The cell count per slide was estimated using the Countess^TM^ Automated cell counter and by counting a fraction of DAPI positive cells on a slide and extrapolation. Zero to few cells with signals for both antibodies were identified on each slide, leading to positive cell ratios of 1 in 10,000 or lower (Table 3, Figure 6). In one sample, two cells positive for *ADAM12* but negative for *HLA-G* were identified as well. Furthermore, there were no cells with *HLA-G*, *MCAM*, or *PSG2* signals only. Sometimes, immunofluorescent signals were present but cells were smaller and had a higher nucleus-to-cytoplasm ratio compared to other positive cells. These cells were classified as ambiguous. In conclusion, our results support the presence of cervical trophoblasts, but at a lower ratio than previously reported, rather in an order of magnitude of 1 in 10,000 than of 1 in 2,000.

**Fig. 6.**
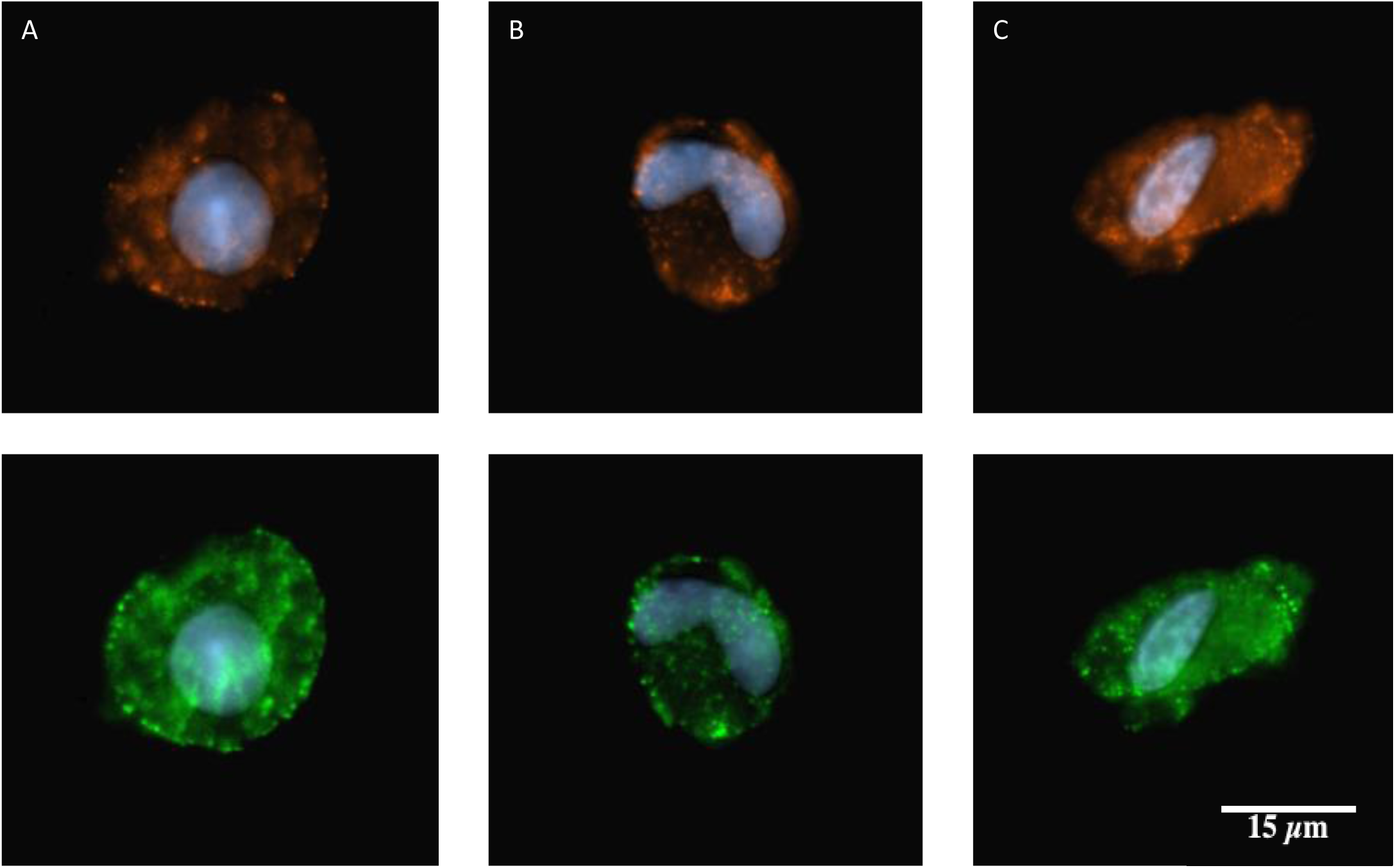
HLA-G positive cell types. Examples of positive cells identified in cervical samples labeled with *HLA-G* (red) and (a) *ADAM12*, (b) *MCAM* and (c) *PSG2*.

**Table 3:** Screening results of stained cervical samples from pregnant women.

| Sample | Antibodies | # cells with double staining per slide | # ambiguous cells with double staining per slide | Estimated cell count per slide |
| --- | --- | --- | --- | --- |
| CC132 | HLA-G + MCAM | 2 | 1 | 65,000 - 90,000 |
| CC133 | HLA-G + ADAM12 | 4 | 2 | 45,000 - 70,000 |
|  | HLA-G + PSG2 | 0 | 0 |  |
| CC134 | HLA-G + ADAM12 | 5 | 3 | 65,000 - 100,000 |
|  | HLA-G + MCAM | 4 | 0 |  |
| CC135 | HLA-G + PSG2 | 2 | 0 | 35,000 - 40,000 |

## Discussion

We present a first single-cell RNA-seq atlas of exocervical smears during early gestation. While recent investigations explored the composition of cervical biopsies during pregnancy in human and mice at the single-cell level^42,43^, our cervical samples are obtained non-invasively. This has as limitation that the number of cells is lower as compared to atlases derived from invasive sampling. We identified two main populations of epithelial cells, as well as various immune cell types. In parallel, a cell atlas of the early placenta expands the existing compendium of trophoblast and mesenchymal core cell subpopulations, and genetic analysis allows for the distinction of cells with maternal and fetal origins. We identified a new subpopulation of EVTs with high expression of epithelial markers. Integration of both cell atlases revealed similarities in the molecular profiles of placental and exocervical cell types. The latter may complicate the identification and isolation of trophoblasts in the exocervix for prenatal testing. In addition, we identified new potential markers that could be used to leverage cervical trophoblast enrichment. Immunocytochemistry shows specificity of these markers for cervical trophoblasts, and indicates their frequency is lower than previously estimated.

The two main types of epithelial cells in the exocervical smears are the squamous epithelial cells that reside at the exocervix and glandular epithelial cells derived from the endocervix. Neutrophils were the most prevalent immune cell type, followed by monocytes/macrophages, T-cells, plasma cells, NK-cells and finally B-cells, with a small population of plasmacytoid dendritic cells. This atlas confirms earlier observations from cytology and flow cytometry: both exo- and endocervical cells are identified in cervical smears, and neutrophils were reported as the most abundant immune cell subtype in cervical smears, both in non-pregnant and pregnant women^18,22,45,46^. Samples from the cervicovaginal interface of women at risk of preterm delivery during early and mid-pregnancy mainly contained neutrophils, and also here variable levels of NK cells, B-cells, T-cells and monocytes were detected^19^. With 1,390 epithelial cells and 9,139 immune cells, the immune cells are by far the largest fraction in our dataset. This conflicts with cytology reports, where epithelial cells were the dominant cell type in cervical smears from pregnant women^19^. Also microscopically, epithelial cells appeared to be more prevalent in our samples (Supplementary Fig. 10). It was noted that epithelial cells are large, deformable and tend to stick to other cells. This makes it difficult to keep them in suspension and capture on the 10x microfluidic system. Therefore, it is possible that the smaller immune cells are overrepresented relative to the larger epithelial cells. Additionally, these cellular characteristics of epithelial cells could complicate the detection and isolation of minor cell subpopulations, such as cervical trophoblasts.

Previous placental single-cell RNA-seq atlases reported heterogeneous cell populations in both early and term, and normal and preeclamptic pregnancies^20,21,28,41,46^. Liu *et al*. and Suryawanshi *et al*. both analyzed first-trimester placental material, between six and eleven weeks gestational age^20,21^. Surprisingly, we detected a novel cluster of EVTs with high expression of epithelial markers. While *HLA-G* expression was limited in this cluster, it was grouped together with other EVT populations and separate from the VCTs. We first hypothesized these EVT cells were precursors of the invasive EVTs, at the beginning of the epithelial-mesenchymal transition (EMT), however, trajectory analysis suggested they arise as a separate group from the VCTs. We further identified one proliferating EVT population, and one with high expression of *SERPINE1*, in accordance with the findings of Liu *et al*. Altogether, the transition reflects the trajectory of EVTs undergoing EMT; EVTs proliferate at the proximal cell columns and further differentiate towards the distal end, where proliferation ceases upon invasion of the decidua and the spiral arteries. In addition, we observe three main and one smaller VCT subtype. One VCT subpopulation expressed *ERVFRD-1*, which are the fusion-competent cells pointing to a pre-SCT state, indicating VCTs function as precursor cells, in accordance with previous observations^21^.

We detected two populations of Hofbauer cells and one population of maternal macrophages. In contrast, Liu *et al*. described two populations of Hofbauer cells but no maternal macrophages^21^. However, one of the Hofbauer populations was characterized by expression of *LYZ* and *HLA-DR*; markers that were highly expressed by our maternal macrophage population. It is possible that these cells were falsely classified as fetal (Hofbauer) cells. Different subpopulations of placenta-associated maternal macrophages were recently identified^36^. Interestingly, our population corresponds to the most abundant subtype of maternal macrophages, which was described to adhere to the placental surface, rather than being part of the mesenchymal core. It was suggested that these cells are present at sites of damage at the syncytium, where they mediate the repair process^36^. The single population of Hofbauer cells described by Suryawanshi *et al*. corresponds to our largest cluster of Hofbauer cells.

Some observations in our analyses raise questions. Our novel EVT cluster identified is composed of cells derived from all samples, making it unlikely to concern a technical artefact, and its markers are also detected in the dataset of Vento-Tormo et al., (Supplementary Data). However, there are some clusters that are dominated by a single sample (clusters 17, 18 and 19 in the placental dataset). Also, we obtain a higher proportion of fibroblasts compared to trophoblasts, conflicting with previous reports^20,28^. It is uncertain whether this is related to the samples or their karyotype, to the sampling method, or whether these are technical artefacts (e.g., doublets).

As mentioned, the frequency of cervical trophoblasts is reported to be 1 in 2,000^37^. As our scRNA-seq dataset consists of 10,539 cells, we expected to identify some trophoblasts. No cells co-expressing established trophoblast marker genes could be identified. However, the immunostaining experiments suggest cervical trophoblasts are present. Immunocytochemistry screening on cervical samples is challenging; the large epithelial cells, especially when sticking together and in the presence of mucus, demonstrate high background staining. For the PSG2 antibody, bright fluorescent signals were detected on a small part of the single cervical epithelial cells as well (Supplementary Fig. 11). Both the fluorescent signals and the cellular morphology of the epithelial cells are different from those in the putative cervical trophoblasts. The frequency of cervical trophoblasts was found to be lower than initially reported^37^. Lower trophoblast frequencies and low yields in trophoblast isolation from cervical smears have now been reported by different teams^11,17,47^. While this could be related to the trophoblast detection and isolation methods, it is possible that trophoblast yield is related to sampling position, with more invasive and potentially more harmful sampling methods obtaining a higher proportion of trophoblast cells^47^.

The few trophoblast isolation methods described today lack reproducibility and are not ready to be clinically implemented^11,39,40,47^. *HLA-G* is the only molecular marker that has been applied for targeting cervical trophoblasts, but seems to be insufficient for pure trophoblast isolation^39,40^. Markers used to isolate trophoblasts from maternal blood fail to isolate EVTs from cervical smears^48^. To identify potential novel markers, we combined both exocervical and placental cell atlases. Surprisingly, co-clustering of placental and cervical epithelial cells was noted. Trophoblasts form the first epithelium during human development and EVTs retain expression of general epithelial markers such as cytokeratin 7 during EMT^13^. The expression of such epithelial markers might confound their isolation or identification. For example, cytokeratin 7 has been used to stain putative cervical EVTs^37^. However, *KRT7* is expressed by cervical epithelial cells as well (Supplementary Fig. 12). Gene expression of other cytokeratin genes (*KRT10*, *KRT18*, *KRT19*) and epithelial markers (*HPGD*, *IFI27*) are shared by invasive EVTs and epithelial cells. Additionally, while it is hypothesized that the invasive EVTs migrate to the cervix, detection of syncytial fragments suggests HLA-G negative trophoblast populations are present as well^11^. To enhance trophoblast isolation, a panel of molecular markers allowing for positive and negative selection might be better suited. We focused on markers unique for trophoblasts or epithelial cells, as immune cells could be more easily removed from the cervical smears using general immune cell markers such as CD45. Our immunocytochemistry analyses suggest these markers could be applied to target cervical trophoblast cells. Further studies combining these markers with isolation techniques such as magnetic- or fluorescence-activated cell sorting are now required to further determine their potential in cervical trophoblast isolation.

In conclusion, we identified all cell types involved in cervical trophoblast isolation, by creating a first cell atlas of exocervical smears obtained from pregnant women. The low cell sample size is a limitation and larger scale sampling across multiple pregnancies may be required to obtain the full compendium of cervical cell types.

Creating cell atlases provides insight into the immune cell types residing at the exocervix during early pregnancy. We envision that this could become a novel approach to study infections and gynecological cancers. Integration of the placental and cervical datasets highlighted the similarity between trophoblasts and maternal epithelial cells, but still allowed for the identification of novel markers specific to both. Further exploration of the differentiating markers between trophoblast and maternal cells could leverage cervical trophoblast isolation.

## Methods

### Sample collection

The studies for collection of exocervical smears (S60410) and placental tissue (S63761) were approved by the Ethics Committee (EC Research) of the University Hospital Leuven. Patients were recruited at the Obstetric Department of the hospital (UZ Leuven, Belgium) and written informed consent was obtained from all participants.

Exocervical smears were collected after confirmation of pregnancy between seven and twelve weeks gestational age. Women with a history of repeated miscarriage, clear signs of vaginal infection or cervicitis, vaginal blood loss during pregnancy or with known infection of human immunodeficiency virus or hepatitis A or B, were excluded from the study. Sampling was carried out using a FLOQSwab^TM^ (Copan Diagnostics) that collects cells at the exocervix after positioning of a speculum. The FLOQSwab^TM^ was then transferred into a 50 mL Falcon tube containing 5 mL of phosphate-buffered saline (PBS). The samples were immediately stored on ice and transferred to the laboratory within a few hours after sampling.

Placental tissue was obtained in parallel with a biopsy for cytogenetic evaluation, in women undergoing a hystero-embryoscopy for a third or more consecutive miscarriage between five and twelve weeks gestational age. Therefore, a biopsy from the chorionic villi was taken under direct hysteroscopic view and wherever possible, the placenta was localized following the umbilical cord to its origin in the inner wall of the chorionic sac. Placental villi were specifically targeted for isolation and removed using a forceps smaller than 2 mm in size.

### Exocervical smear processing

After removing the FLOQSwab^TM^ from the Falcon tube, the side of the tube was washed with 5 mL of PBS to flush remaining cells, after which all cells were collected by centrifugation at 300g for 5 min. An additional washing step was performed, and cells were resuspended in 1% BSA in PBS. In case large cell clumps were present, cells were strained using a 70 µm Flowmi^TM^ cell strainer (Bel-Art SP Scienceware) to avoid clogging. Sample constitution and cell viability were assessed using the LUNA-FL^TM^ Automated Cell Counter (Westburg) (Supplementary Fig. 13).

### Placental tissue processing

Placental tissue processing took place immediately after sampling. If visibly present, maternal material was removed from the chorionic villi using forceps. A small part of the villous biopsy was separated for DNA analysis. The main part of the biopsy was kept for scRNA-seq, for which tissue was dissociated as described by Suryawanshi et al.^20^, with some minor modifications. Filtration was carried out using 70 and/or 40-micron Flowmi^TM^ cell strainers (Bel-Art SP Scienceware), depending on the amount of larger debris. Final resuspension was done in ice-cold 1x PBS with 0.04% BSA and 1U/µL SUPERase-In RNase Inhibitor (Thermo Fisher Scientific). Cell viability, concentration, and the amount of debris were assessed using the LUNA-FL^TM^ Dual Fluorescence Cell Counter (Westburg) following the manufacturer’s instructions (Supplementary Fig. 13).

### Single-cell RNA sequencing and analysis

Single-cell sequencing libraries were generated using the Chromium Single Cell 3’ Reagent Kit (v3.1) (10x Genomics) following the manufacturer’s protocol. Libraries were sequenced on the NovaSeq 6000 (Illumina). Sequencing data were aligned and quantified using the Cell Ranger software (10x Genomics) and the reference index was built using the GRCh38 human reference genome. The filtered matrices generated by Cell Ranger were then read in the Seurat (version 4.2.0) R package^49^. Cells with less than 100 detected genes or less than 500 reads were excluded, as well as cells with total mitochondrial gene expression above 10% (Supplementary Fig. 14). Ambient RNA was removed from exocervical smear data using SoupX (version 1.5.2)^50^. Doublets were flagged and removed using the R package DoubletFinder (version 2.0.3)^51^. Samples were merged in Seurat and normalization and variance stabilization were carried out using the SCTransform function. Mitochondrial gene expression was regressed out during this step for both placental and exocervical data, and for the latter, ribosomal gene expression and expression of heat shock proteins were also regressed out. Then, principal component analysis was done with Seurat and samples were integrated using the R package harmony (version 0.1.0)^52^. Further downstream analyses, including shared nearest neighbor graph-based clustering and differential expression analysis were performed using Seurat^49^. While we aimed to generate gene expression data of 10,000 cells per sample, we systematically obtained lower yields per sample for the exocervical smears (Supplementary Table 1). This is probably related to the relatively low cell viabilities that were obtained, which could be caused by the high turnover rate of the cervical epithelium^53^.

Differentially expressed genes for each cluster were identified using the FindAllMarkers function implemented in Seurat using the Wilcox test with the following parameters: (1) only positive markers were selected, and (2) only genes with a p-value below 0.05, (3) that were expressed in at least 25% of cells (4) with a minimum difference of 0.05% in the fraction of detection between the two groups were considered. To find differentially expressed genes between specific clusters or cell types, the FindMarkers function was used and only positive markers that were expressed in at least 10% of all cells were considered. For the combined analysis in which markers specific to placental or epithelial cells were explored, differentially expressed genes were sorted based on the log fold-change of the average expression between the two groups considered. Only markers with values higher than one were selected. As the corresponding proteins should be accessible by antibodies, further selection was done based on a transmembrane gene list assembled by Dannenfelser et al^54^. Expression of those markers was then explored in the placental and exocervical datasets to assess their specificity, using the FeaturePlot function of Seurat.

Functional signatures of the clusters were generated based on differentially expressed genes using the R package clusterProfiler (version 4.2.2)^55^. The CellCycleScoring function of Seurat was used to classify cells into discrete phases of the cell cycle. Maternal contamination in the placental villous was explored for each sample using the package freemuxlet, which employs single-nucleotide polymorphisms (SNP) information from the scRNA-seq data to deconvolute different genotypes (https://github.com/statgen/popscle). The same tool was used in an attempt to identify fetal (placental) SNPs in exocervical smears. Monocle3 and Slingshot were applied for trajectory analysis of trophoblast cells^56,57^.

### Aneuploidy screening of chorionic villi

To assess the copy number status of the placental material, part of the placental sample was subjected to DNA extraction using the DNeasy Blood and Tissue kit (Qiagen) following the protocol ‘Isolation of Genomic DNA from Tissues’. DNA quality and concentration were assessed by NanoDrop ND-1000 Spectrophotometer (Thermo Fisher Scientific) and Qubit 2.0 Fluorometer (Invitrogen). Analysis of the copy number profile was then carried out by array comparative genomic hybridization (aCGH) using the 8×60 K CytoSure ISCA v3 microarray (Oxford Gene Technology). Results were analyzed using the CytoSure Interpret software (Oxford Gene Technology).

### Immunocytochemistry

Leftover CVS samples were used to test reactivity of the selected antibodies for trophoblast cells. CVS samples were dissociated to obtain a single-cell suspension by resuspension in 1/6 collagenase (CLS II, Worthington Biochemical Corporation) diluted in DMEM/12 culture medium (Gibco, Thermo Fisher Scientific) and overnight incubation at 37°C. The next day, the sample was centrifuged at 300g for 10 min and the supernatant was removed. For immunofluorescent staining of cervical samples, the FLOQSwab^TM^ was rinsed in the 50 mL tube containing 5 mL of PBS. An additional 5 mL of PBS was added to collect remaining cells from the side of the tube. All liquid was then transferred to a 15 mL tube and centrifugation was carried out at 300g for 10 min. Immunostaining on both cervical and CVS samples was carried out as follows: cells were first fixed in 2% PFA for 10 min at room temperature, followed by a washing step in 3 mL of PBS and centrifugation as described. The supernatant was removed and cells were resuspended in BlockAid blocking solution (Invitrogen) and incubated on ice for 1 hour. Then, primary antibodies where added and incubated for 1 hour on ice. Subsequently, cells were washed in PBS as described above, and resuspended in secondary antibody in a final concentration of 1 in 600 and incubated for 1 hour on ice, followed by another washing step. The different antibodies and their final concentrations are listed in Supplementary Table 10. Final resuspension was done in PBS. Slides for microscopic analysis were prepared by adding 10 µL of cell suspension on a slide with an equal amount of VECTASHIELD HardSet Antifade Mounting Medium with DAPI (Vector Laboratories) to visualize nuclei. A coverslip was immediately put on top. Slides were cured for 15 min at room temperature and subsequently stored at 4°C before microscopic evaluation. Visualization was carried out using an epifluorescence microscope (Zeiss Axioplan) using Cytovision software (Leica Biosystems).

## Declaration statements

### Data Availability Statement

All sequencing data have been submitted to the European-Genome-phenome Archive (https://ega-archive.org/) under accession number EGAS00001007044.

## Acknowledgements

This research has received funding from the Research Foundation Flanders (FWO) (1S10119N to M.v.R. and 1241121N to O.T.) and from the European Union’s Horizon 2020 research and innovation program under grant agreement No 824110 – EASI-Genomics (J.R.V.). Institutional support was received from the KU Leuven, C1-C14/18/092 and C14/22/125 to J.R.V. We are grateful to all patients who participated in this study. We thank the Genomics Core team for their help with the experiments, the VIB Single Cell Core for running the freemuxlet software, and Kate Stanley for a critical reading of the manuscript.

## Author Contributions

M.v.R. designed and conceptualized the study; collected, analyzed and interpreted the data and drafted/revised the manuscript. J.R.V. and O.T. designed and conceptualized the study and drafted/revised the manuscript. I.P. collected the aCGH data and revised the manuscript. L.L., A.P. and D.T. collected the samples and revised the manuscript.

