## Supplementary Data for "Single-cell dissection of cervix and placenta reveal both novel and overlapping cell types"

**Supplementary Data:
Comparison of our recurrent miscarriage dataset with the dataset from elective pregnancy terminations of Vento-Tormo *et al.***^1^

All maternal cell populations were removed from the dataset of Vento-Tormo *et al.*, leaving 17,443 placental cells. As opposed to our in-house dataset which mainly consists out of aneuploid placental samples, whole-genome sequencing of the placental samples of Vento-Tormo et al. indicated only one fetal genetic aberration which was a trisomy 18. In a first round of analysis, all mitochondrial genes were removed from our dataset as this was done in the dataset of Vento-Tormo as well. Then, we noted differences in gene names in the dataset of Vento-Tormo compared to our dataset, which falsely came out as differentially expressed. Therefore, only genes present in both datasets were retained. UMAPs of the integrated dataset show that the cell annotations as given by Vento-Tormo *et al*. coincide with our own cluster annotations (Figure 2). There is an overlap of VCTs, EVTs, SCTs, fibroblast populations (fFB/FB), Hofbauer cells (HB/HC) and vascular endothelial cells (Endo (f)/VEC). Maternal macrophages are not present in the latter, as we preselected fetal cell populations only. In accordance with Vento-Tormo *et al*., there is a separate cluster of fibroblast cells, annotated by Vento-Tormo *et al*. as fFB2. Marker genes of fFB2 are *PLA2G2A*, *AREG* and *REN*, corresponding to cluster 14 in our own placental dataset analysis. While the same cell populations are present, there are differences in the number of cells contributing to each cell type.

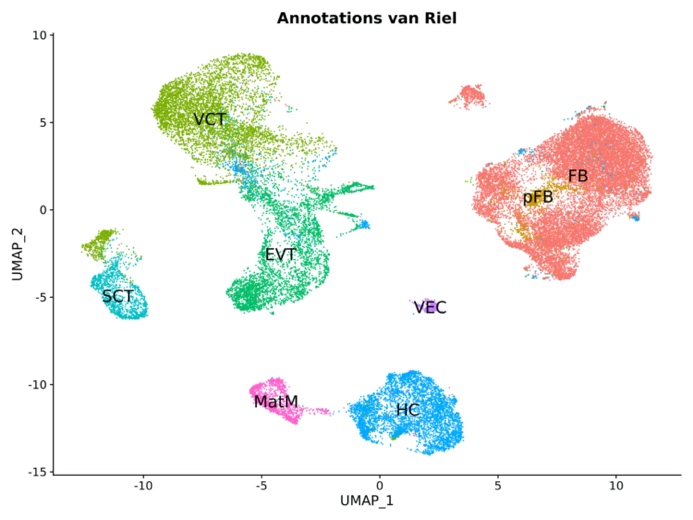

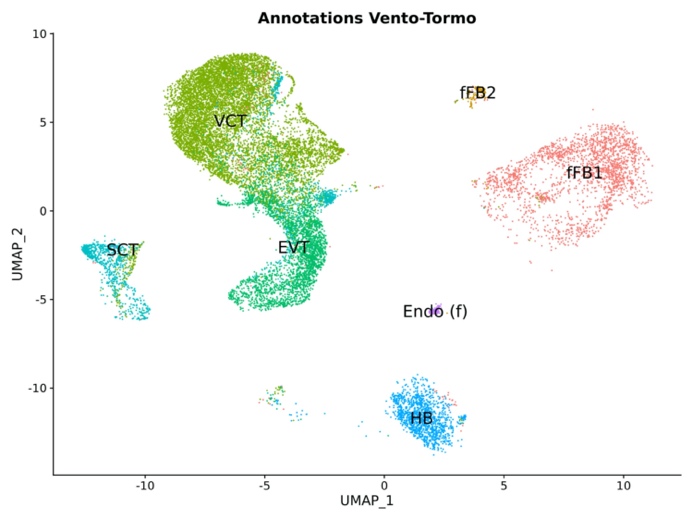

a

b

Supplemental Data Figure 1. UMAPs of the integrated dataset consisting of our data and the placental cells of Vento-Tormo *et al.*, with (a) label transfer of the annotations given by Vento-Tormo et al., and (b) our own annotations.

Supplemental Data Figure 2a shows the UMAP split over the two datasets; while our dataset contains more fibroblast cells (Clusters 1, 2, 8, 12 and 15), the one from Vento-Tormo *et al*. contains more VCTs (clusters 0 and 6). Fibroblast cluster 15 is unique for our dataset, moreover, it only consists out of cells derived from sample PLAC09.

Supplemental Data Figure 2: Clustering results of our dataset and the one of Vento-Tormo *et al*. a. UMAP of the integrated dataset split by the two different datasets. b. UMAP showing the different clusters of the integrated dataset of our samples and those of Vento-Tormo *et al*. c. Bar plot showing the relative frequency of the contribution of each dataset to the different clusters.

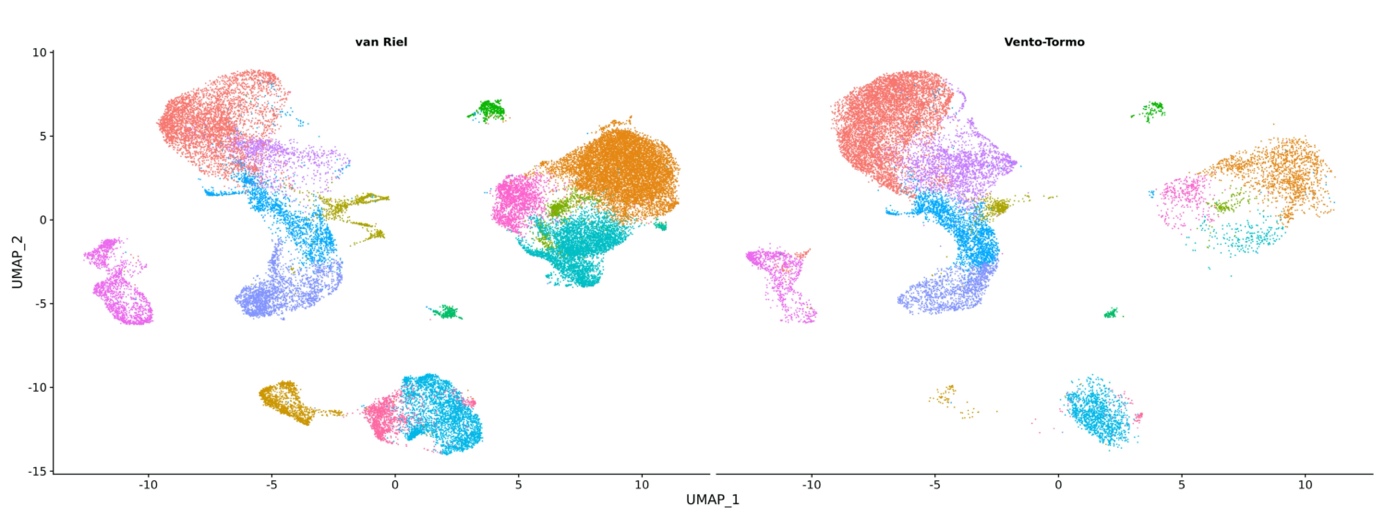

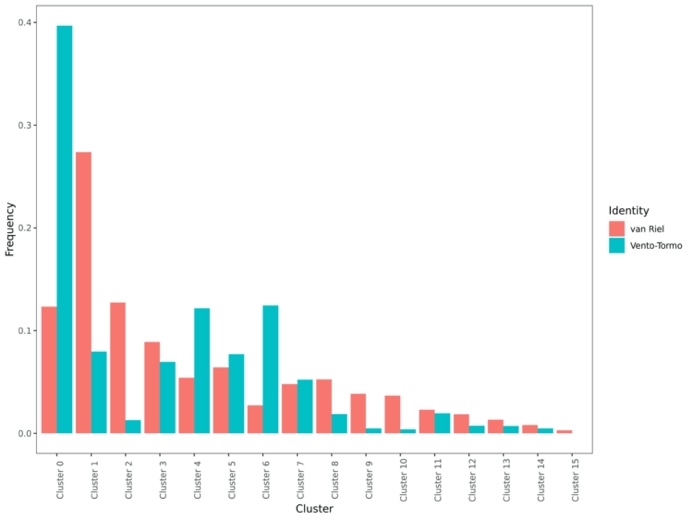

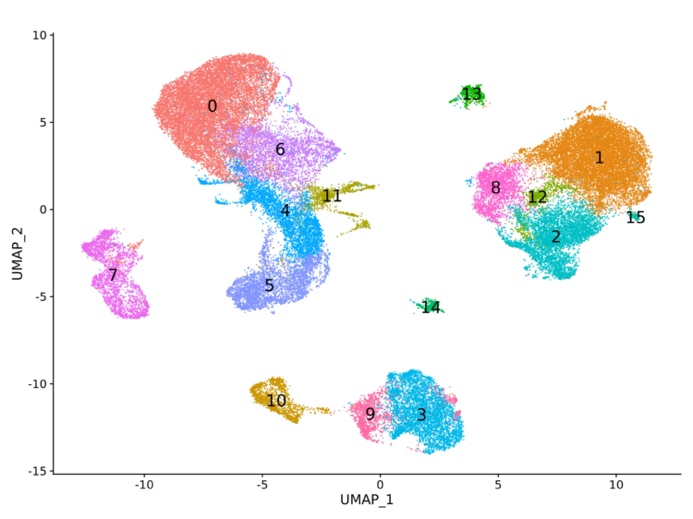

a

b

c

While almost all Hofbauer cells are part of cluster 3 for the dataset of Vento-Tormo et al., Hofbauer cells in our dataset are divided over cluster 3 and cluster 9. When comparing clusters 9 and 3, cluster 9 has upregulation of *CXCL14*, *IGFBP3* and collagen genes, corresponding to cluster 6 in our own placental dataset. Cluster 3 on the other hand has upregulation of *CD14*, *CCL3* and *CCL4*, as described for Hofbauer cells^2^. It is unclear whether the lack of expression of these markers in cluster 10 is related to the fact that these samples are derived from (recurrent) miscarriages.

To validate the marker genes used and identified in our analyses, we subset the dataset of Vento-Tormo and explored gene expression. Expression of the markers used in our dataset to distinguish different cell populations (Figure 3b) and different EVT clusters (Figure 4) matches with the expression in the dataset of Vento-Tormo et al., (Supplemental Data Figure 3 and 4). The fraction of cells expressing GABRP, the marker genes of the EVT cluster with an epithelial phenotype, is however small in the dataset of Vento-Tormo et al.

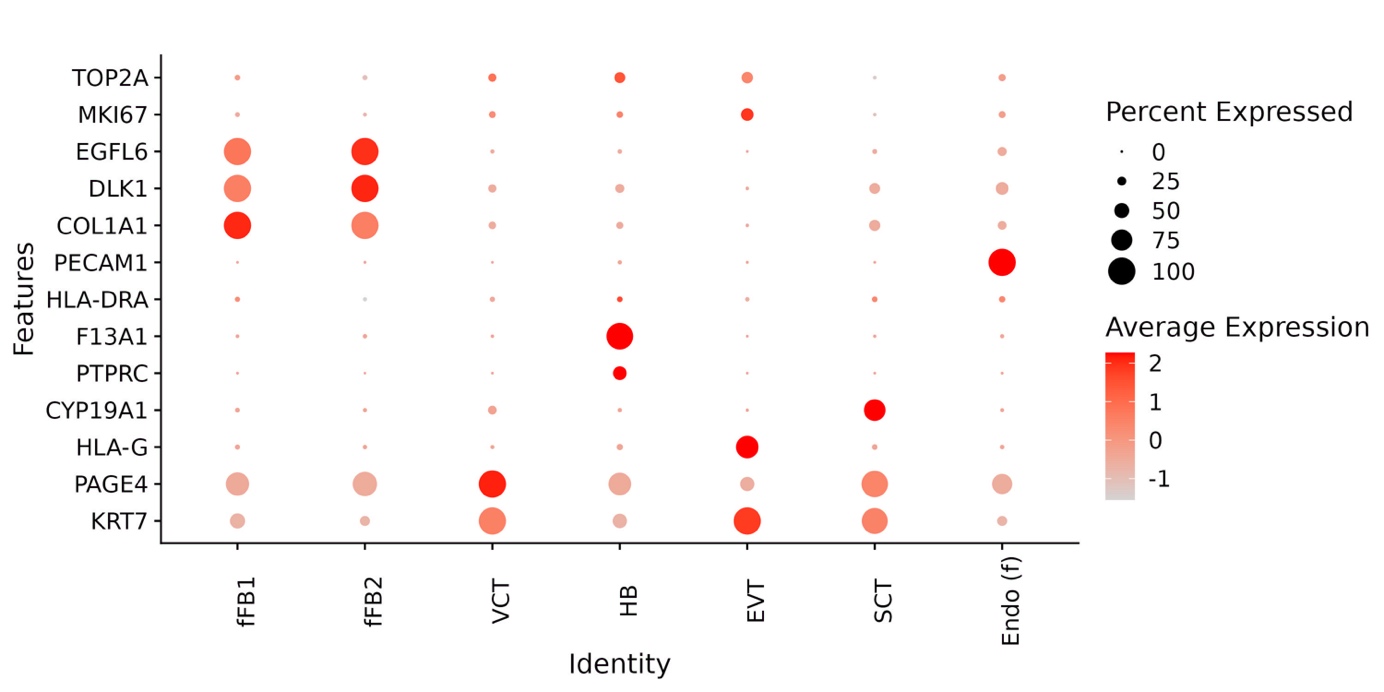

Supplemental Data Figure 3: Dot plot showing expression of established marker genes per cell type as annotated by Vento-Tormo et al.

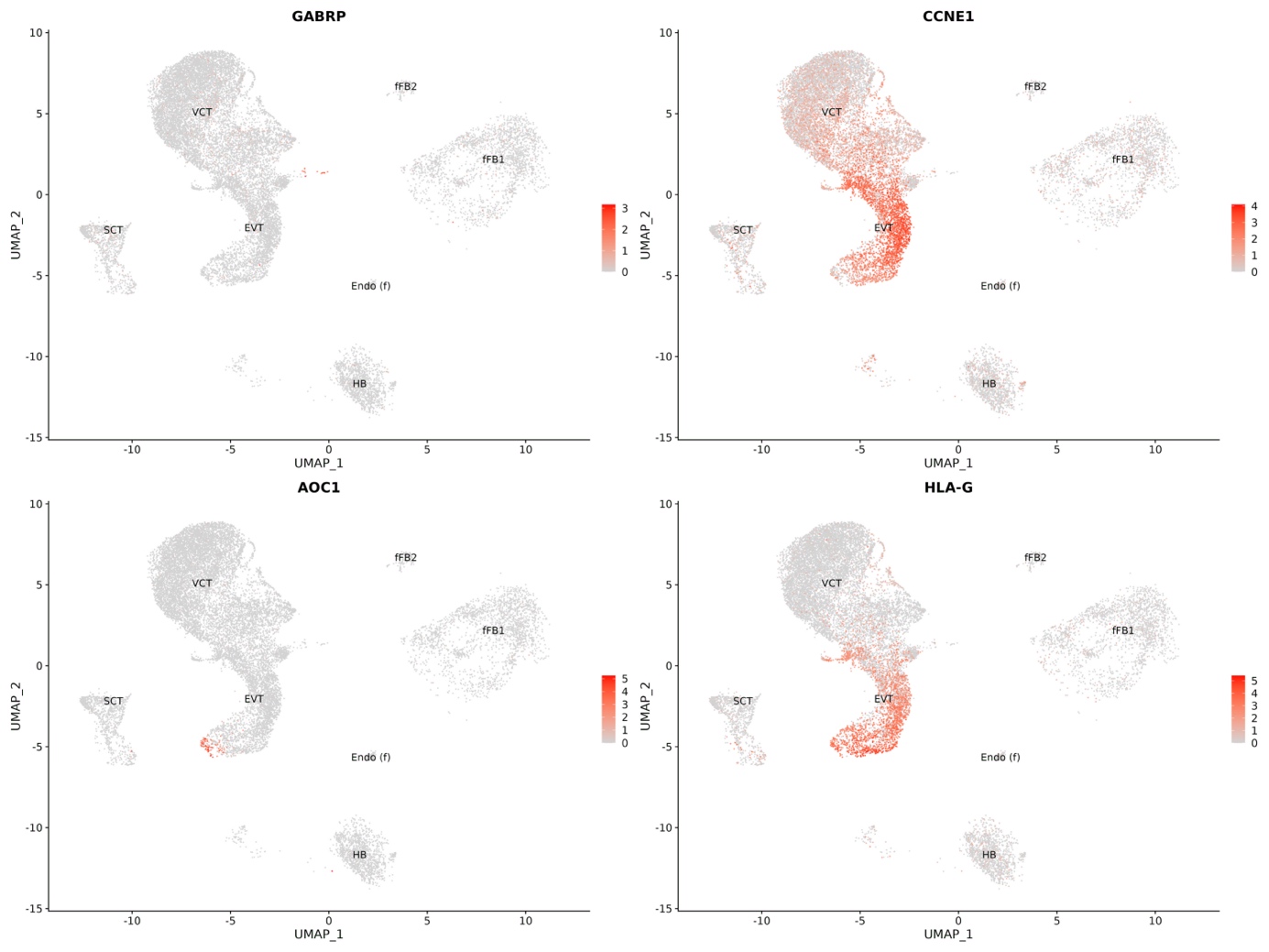
Supplemental Data Figure 4: Feature plots showing expression of the same markers distinguishing the different EVT subtypes in our placental dataset. Similar to our dataset, there is a small number of cells expressing *GABRP*, marker gene of the EVT subtype with an epithelial phenotype. Furthermore, there are proliferating cells, as characterized by *CCNE1* expression, and cells expressing *HLA-G* and *AOC1*, markers of final differentiated EVTs.

In addition, we further explored the newly identified EVT cluster characterized by *GABRP* expression. Other marker genes are also present in the dataset of Vento-Tormo et al (Supplemental Data Figure 5). In the combined dataset, this cluster is cluster 12, consisting of 991 cells, of which 65% is derived from our dataset and 35% from the dataset of Vento-Tormo et al. In addition, there are two samples in each dataset that constitute the largest part of the cluster (55% combined), but all samples contribute, suggesting no batch effect (Supplemental Data Figure 5 and 6, Supplemental Data Table 1).

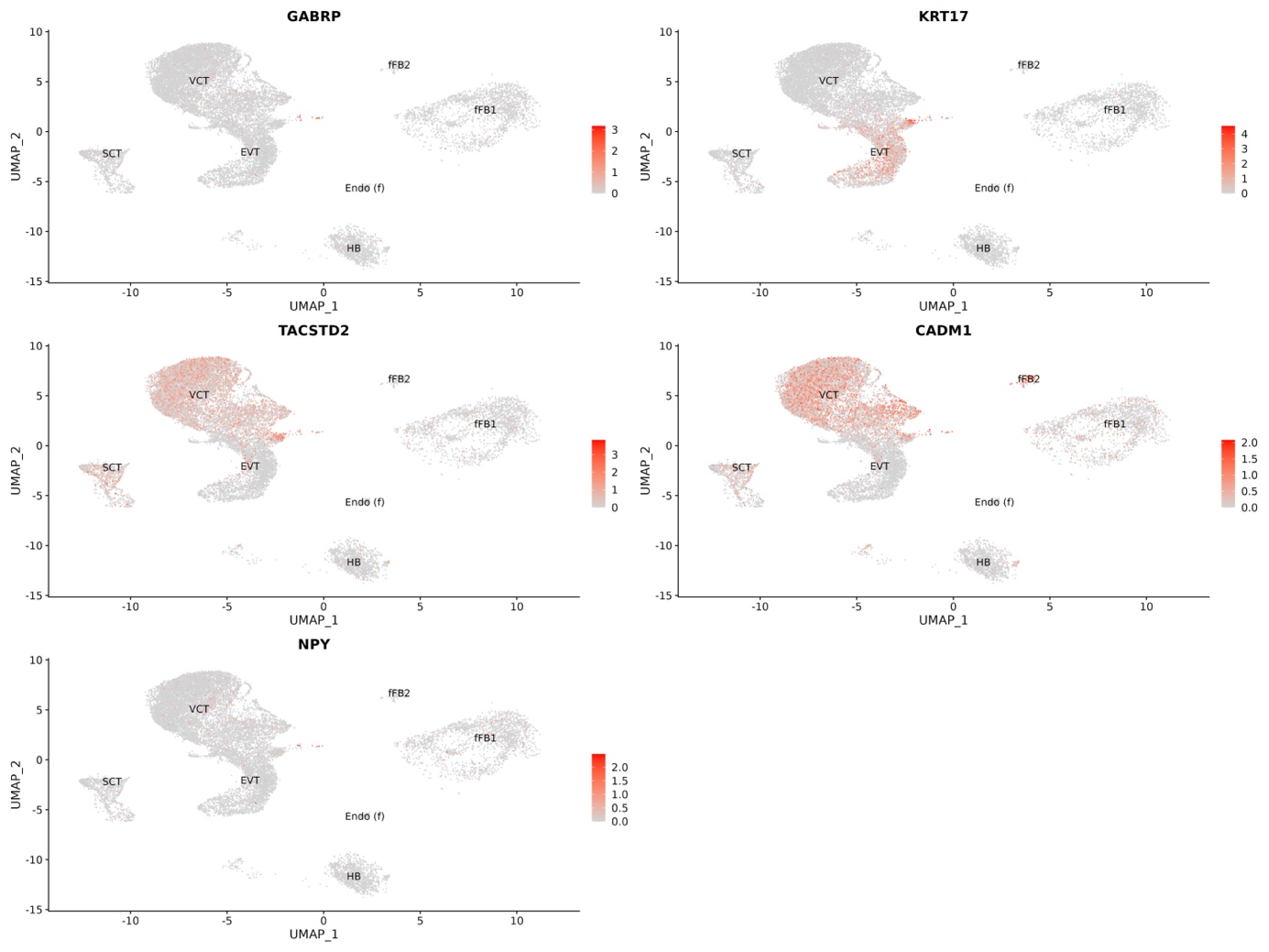

Supplemental Data Figure 5: Feature plots of the marker genes of the placental cells with an epithelial phenotype, plotted on the dataset of Vento-Tormo et al.

Supplemental Data Figure 6: a. UMAP of the integrated dataset. b. Feature plot showing expression of *GABRP*. c. UMAP showing the sample origin plotted separately for each dataset.

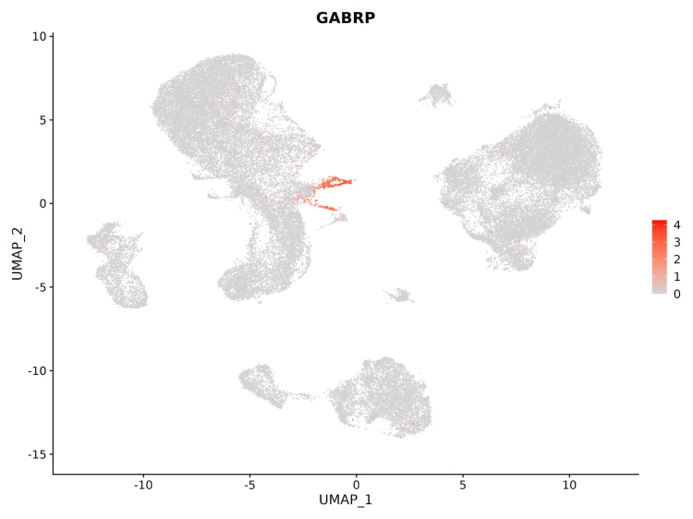

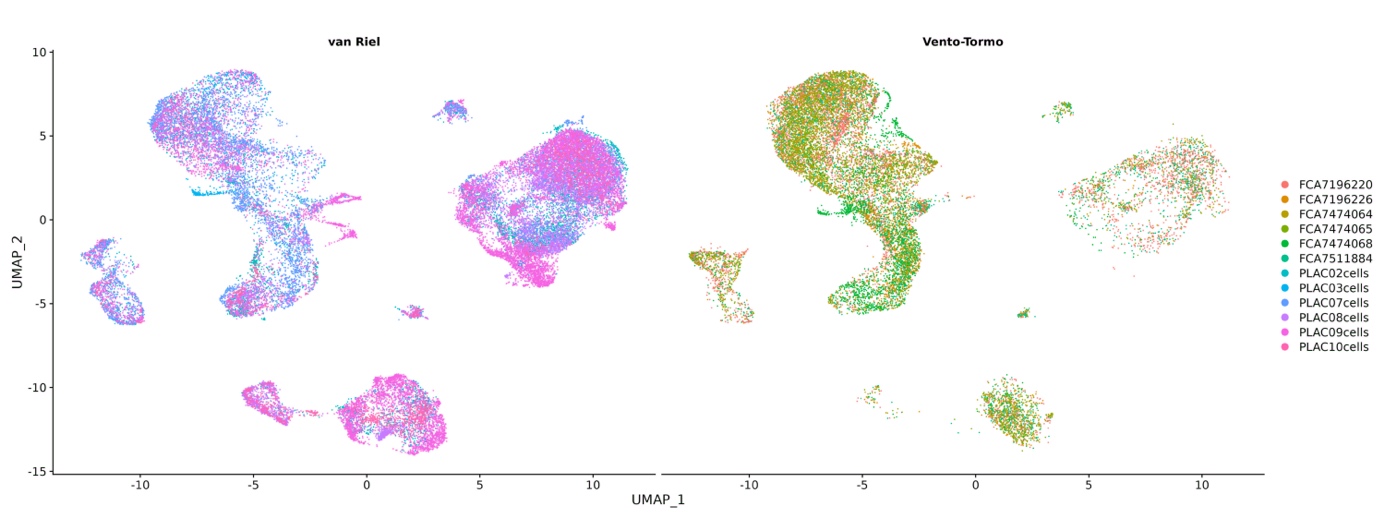

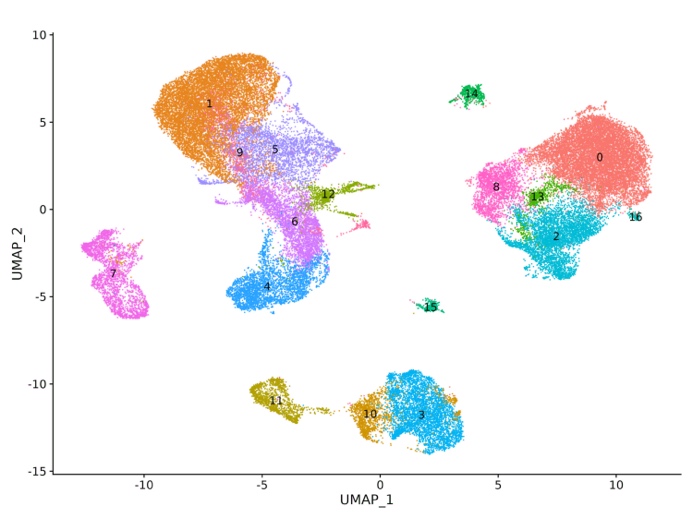

| **Dataset** | **Samples** | **Clusters** | | | | | | | | | | | | | | | | |
| --- | --- | --- | --- | --- | --- | --- | --- | --- | --- | --- | --- | --- | --- | --- | --- | --- | --- | --- |
|  |  | **0** | **1** | **2** | **3** | **4** | **5** | **6** | **7** | **8** | **9** | **10** | **11** | **12** | **13** | **14** | **15** | **16** |
| **Vento-Tormo^1^** | **FCA7196220** | **630** | **1966** | **116** | **162** | **72** | **459** | **279** | **333** | **126** | **50** | **25** | **10** | **100** | **44** | **9** | **12** | **0** |
|  | **FCA7196226** | **84** | **171** | **7** | **140** | **270** | **107** | **461** | **50** | **15** | **5** | **12** | **13** | **38** | **14** | **1** | **7** | **0** |
|  | **FCA7474064** | **139** | **2061** | **18** | **285** | **83** | **542** | **197** | **197** | **33** | **21** | **15** | **13** | **33** | **14** | **33** | **9** | **0** |
|  | **FCA7474065** | **150** | **1942** | **12** | **294** | **77** | **487** | **221** | **198** | **37** | **16** | **16** | **10** | **29** | **19** | **29** | **14** | **0** |
|  | **FCA7474068** | **73** | **84** | **17** | **123** | **607** | **323** | **917** | **93** | **71** | **20** | **11** | **10** | **78** | **7** | **7** | **2** | **0** |
|  | **FCA7511884** | **305** | **615** | **56** | **209** | **82** | **193** | **169** | **69** | **52** | **21** | **6** | **9** | **69** | **30** | **46** | **37** | **0** |
| **van Riel** | **PLAC02cells** | **1257** | **527** | **923** | **307** | **590** | **135** | **104** | **217** | **296** | **245** | **210** | **94** | **36** | **72** | **117** | **51** | **0** |
|  | **PLAC03cells** | **435** | **229** | **66** | **126** | **4** | **197** | **70** | **53** | **39** | **56** | **53** | **19** | **6** | **34** | **18** | **13** | **0** |
|  | **PLAC07cells** | **870** | **1118** | **528** | **18** | **540** | **471** | **568** | **793** | **162** | **851** | **21** | **35** | **96** | **91** | **186** | **43** | **0** |
|  | **PLAC08cells** | **3792** | **413** | **1134** | **576** | **551** | **152** | **96** | **170** | **589** | **309** | **21** | **606** | **48** | **146** | **67** | **40** | **0** |
|  | **PLAC09cells** | **2465** | **815** | **1702** | **1601** | **217** | **139** | **26** | **361** | **760** | **440** | **854** | **253** | **443** | **287** | **68** | **85** | **102** |
|  | **PLAC10cells** | **533** | **151** | **76** | **443** | **242** | **39** | **76** | **84** | **40** | **54** | **168** | **261** | **15** | **15** | **4** | **46** | **0** |

Supplemental Data Table 1: Quantification of the number of cells from each sample contributing to the different clusters.

**Supplementary Figures**

Supplementary Figure 1: Feature plots of markers specific for plasmacytoid dendritic cells (*LILRA4*, *IRF7*, *PACSIN1* and *TCF4*) and B-cells (*MS4A1* and *BANK1*). Cluster 13 (marked with a rectangle) is divided in two groups of cells of which the left part expressed plasmacytoid dendritic cell markers and the right part B-cell markers.

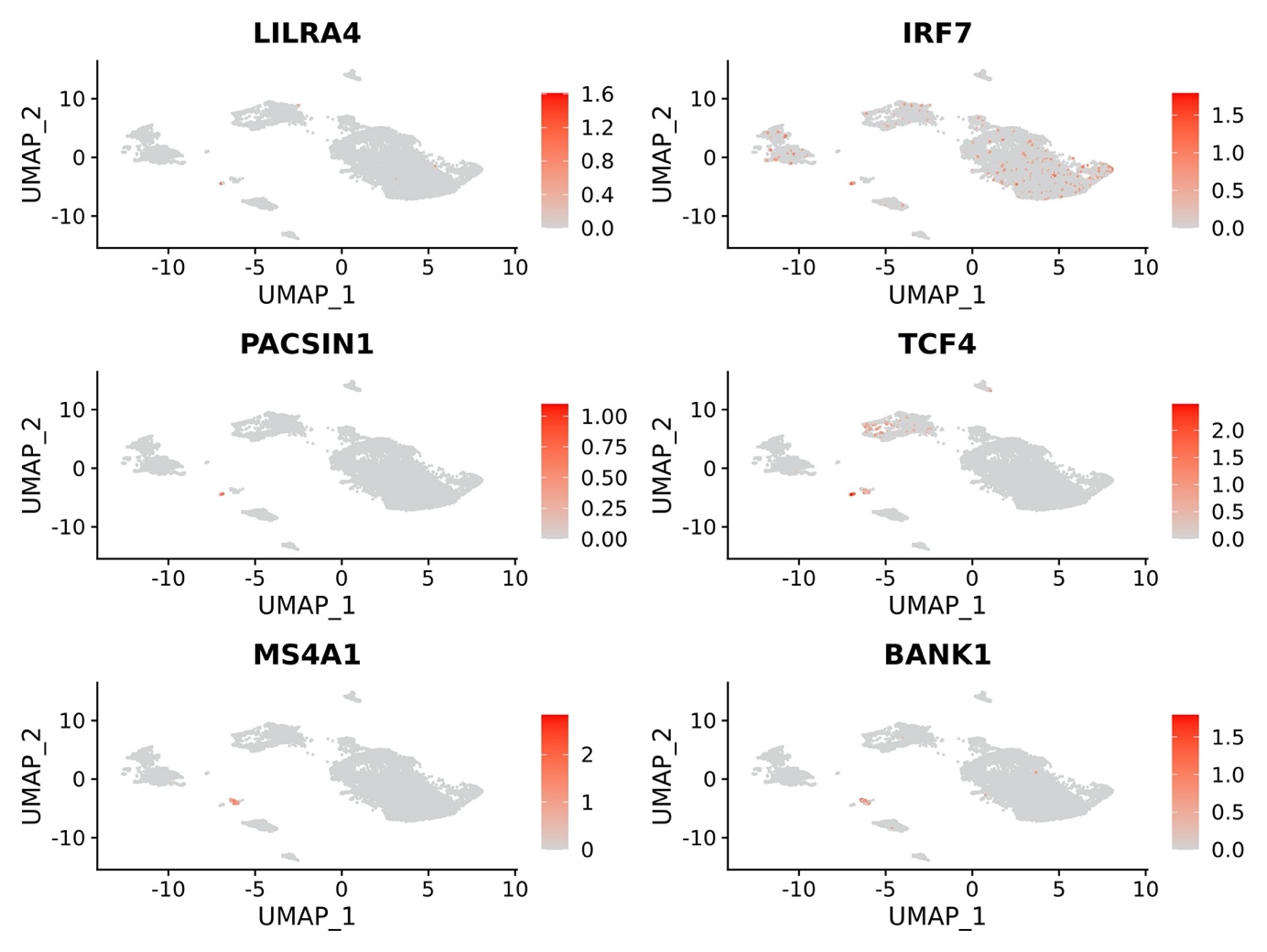

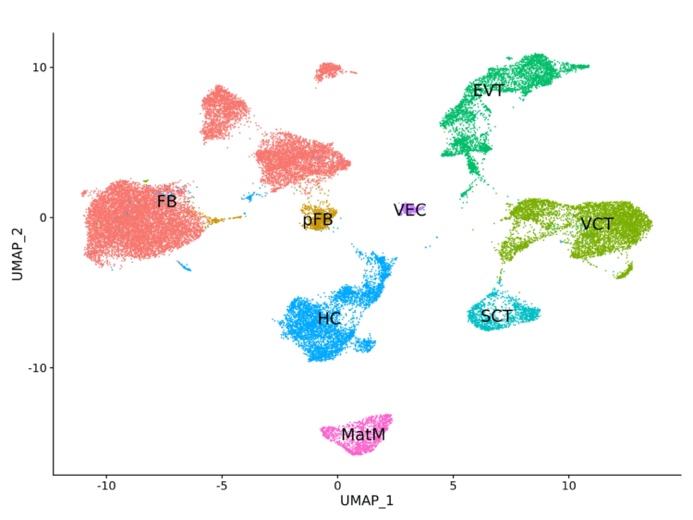

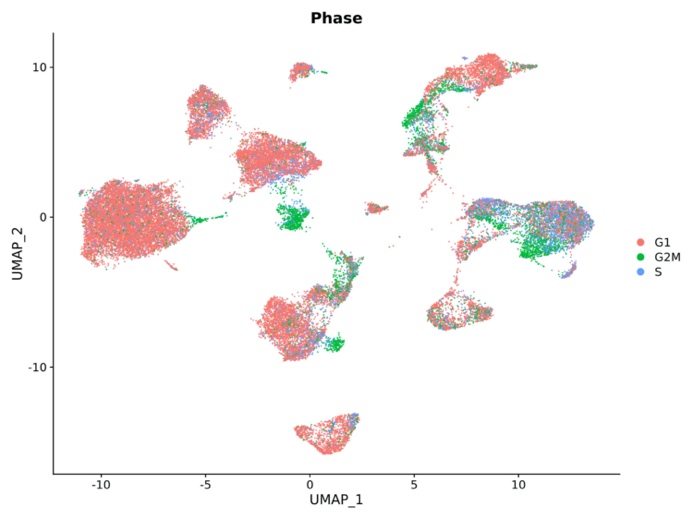

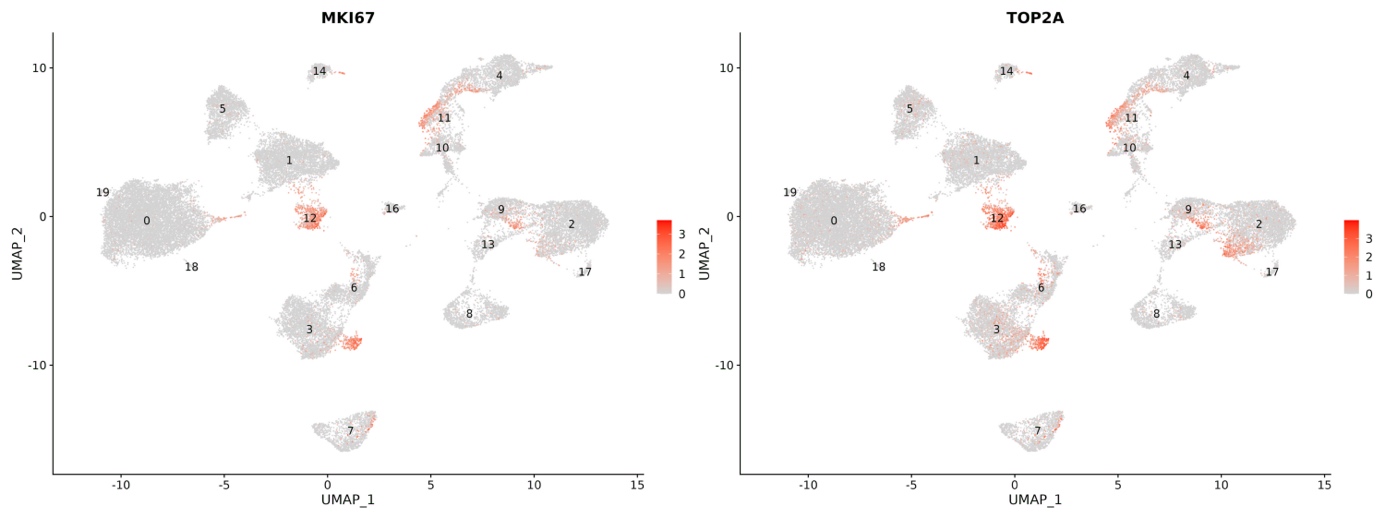
 Supplementary Figure 2: Cell proliferation in the placental dataset. a. Feature plots of cell proliferation markers *MKI67* and *TOP2A*. b. UMAP plot of the cell cycle phase of each cell as determined by Seurat. c. UMAP plot with annotated cell types.

c

a

b

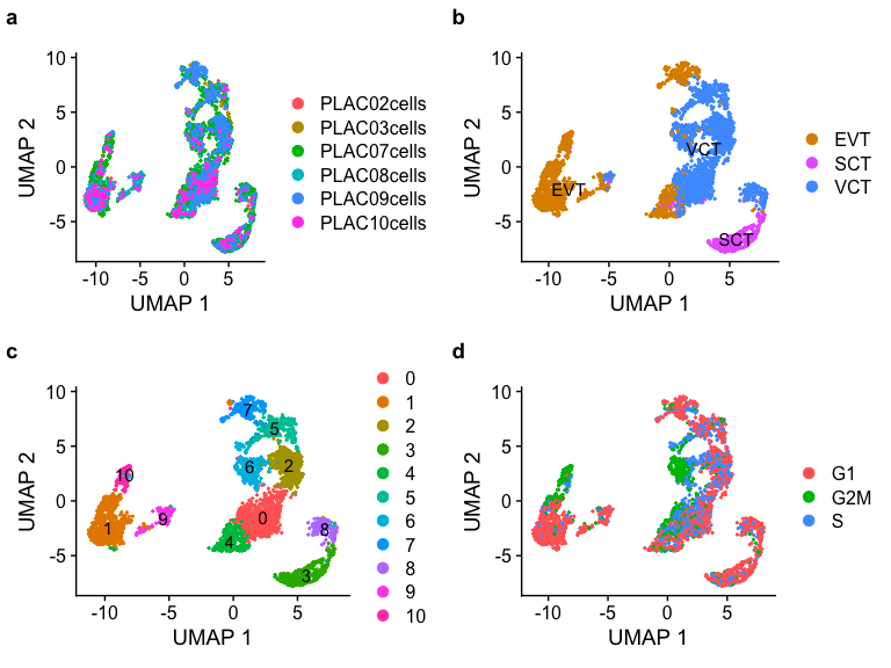
Supplementary Figure 3: Results of the trajectory analysis. a. UMAPs after subclustering of all trophoblast cells showing the sample contributions and cell type annotations. b. Trajectory analysis using Monocle 3. In parallel, trajectory analysis was done using Slingshot, which gave similar results (results not shown).

b

a

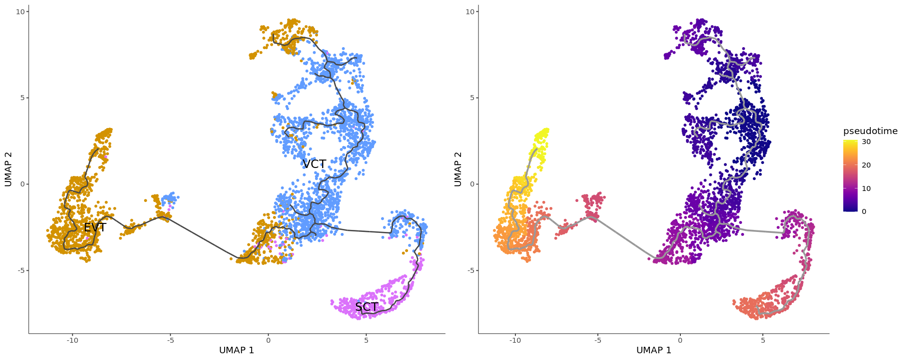

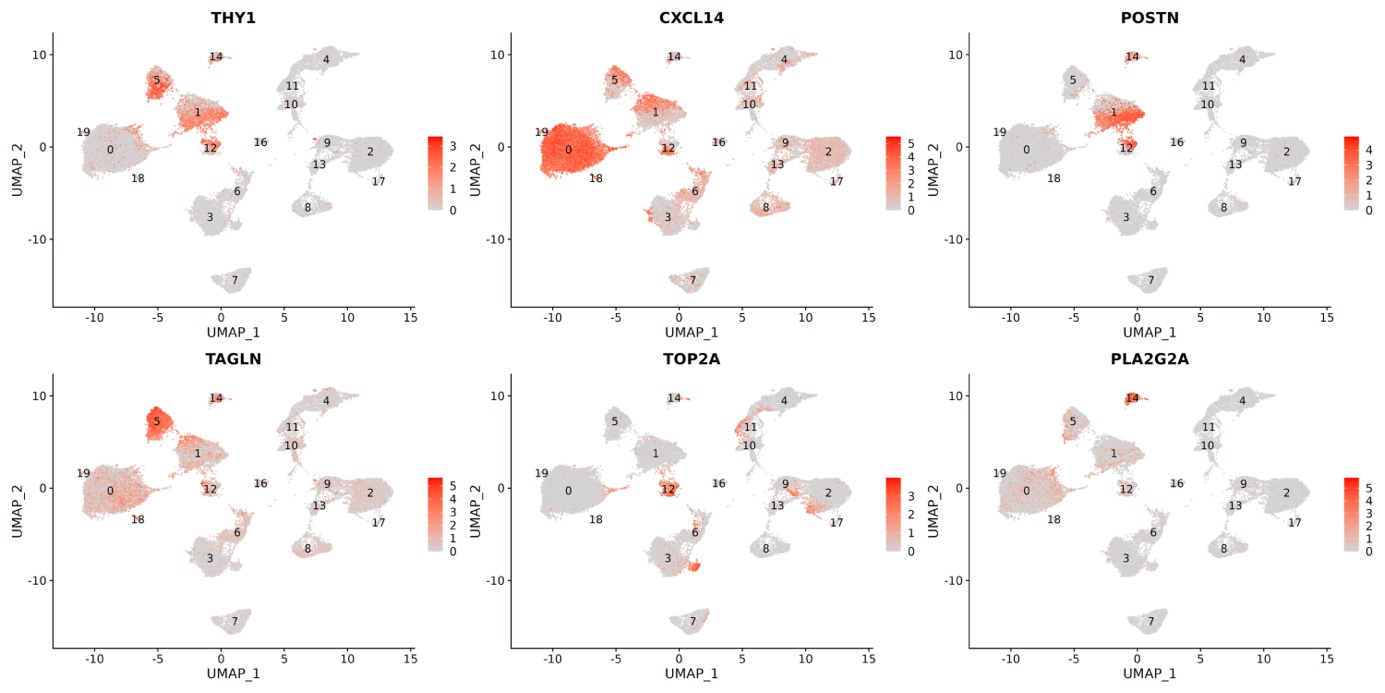
Supplementary Figure 4: Differentially expressed genes between the different main fibroblast clusters (cfr Table 2). Cluster 0 has reduced expression of *THY1* and the highest expression of *CXCL14*. Cluster 1 and 5 are characterized by high expression of *POSTN* and *TAGLN*, respectively. Cluster 12 has high expression of proliferation marker *TOP2A* and cluster 14 is characterized by high expression of *PLA2G2A*.

Supplementary Figure 5: (a) Feature plots showing high expression of both *CCL3* and *CCL4* in Hofbauer cell cluster 3 compared to Hofbauer cell cluster 6. (b) UMAP highlighting cells of cluster 3 (red) and cluster 6 (blue).

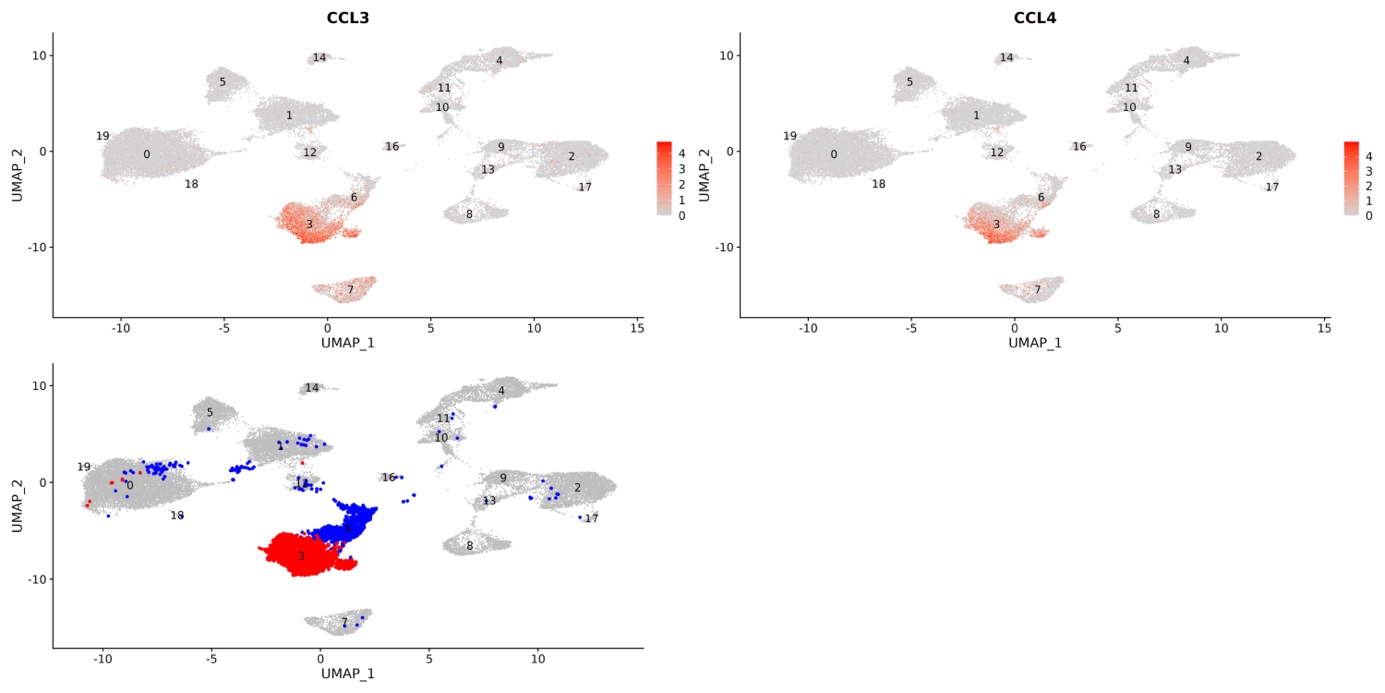

a

b

Supplementary Figure 6: Comparison of markers of Hofbauer cells (clusters 3 and 6) and maternal macrophages (cluster 7). Both express *SPP1*, while *RNASE1* and *CD14* are only moderately expressed by maternal macrophages, and *F13A1* and *LYVE1* expression is absent. Markers of maternal macrophages include *HLA-DRA*, *APOC1* and *LYZ*.

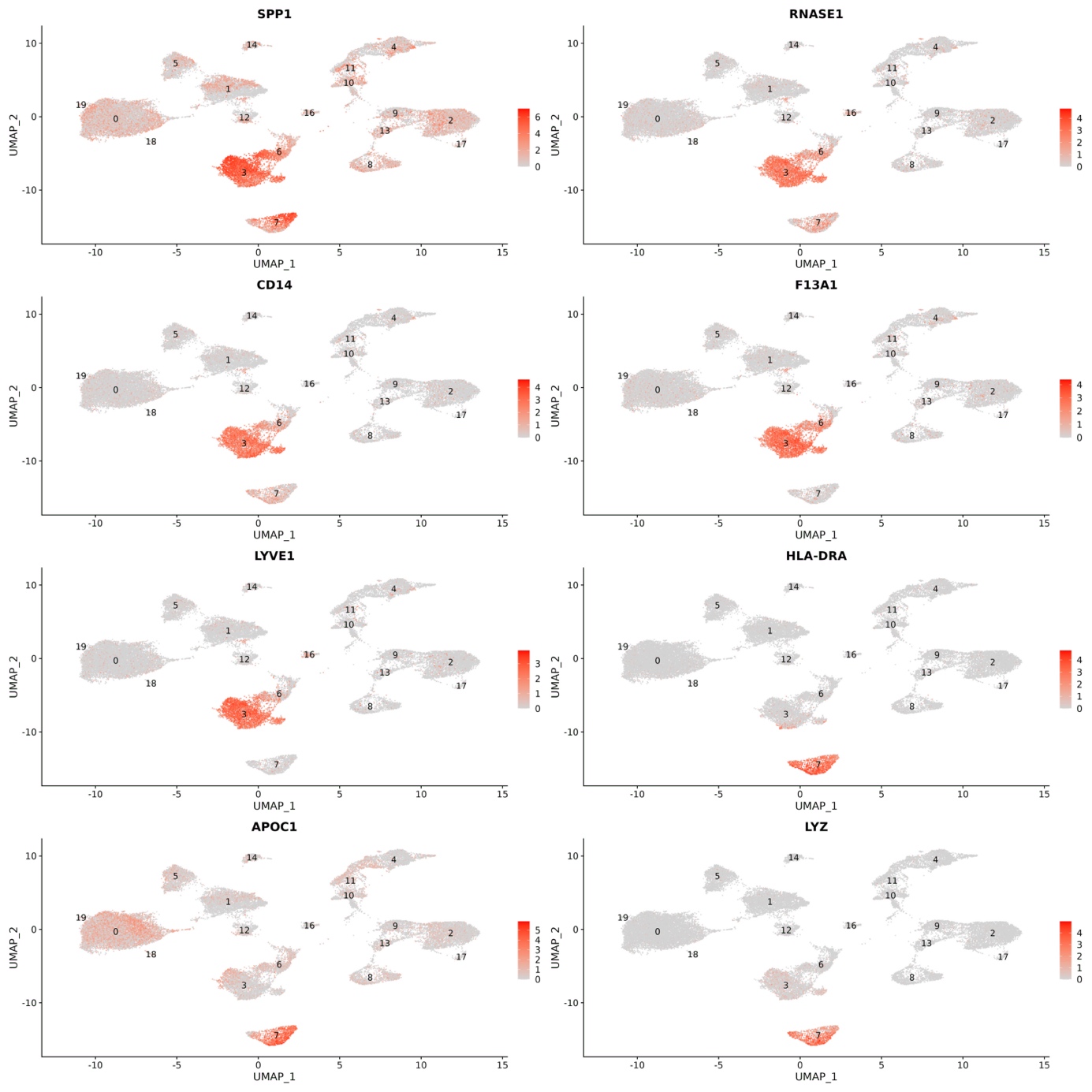

Supplementary Figure 7: Freemuxlet results of the cervical samples. Different cells were labeled as having a different genotype (1,1) from the main background (0,0), but they did not cluster together. Additionally, the parameters reflecting the reliability of the freemuxlet results are better for the placental than the exocervical dataset (data not shown), which is probably caused by the lower cell viabilities in the latter, making the freemuxlet results unreliable in these samples.

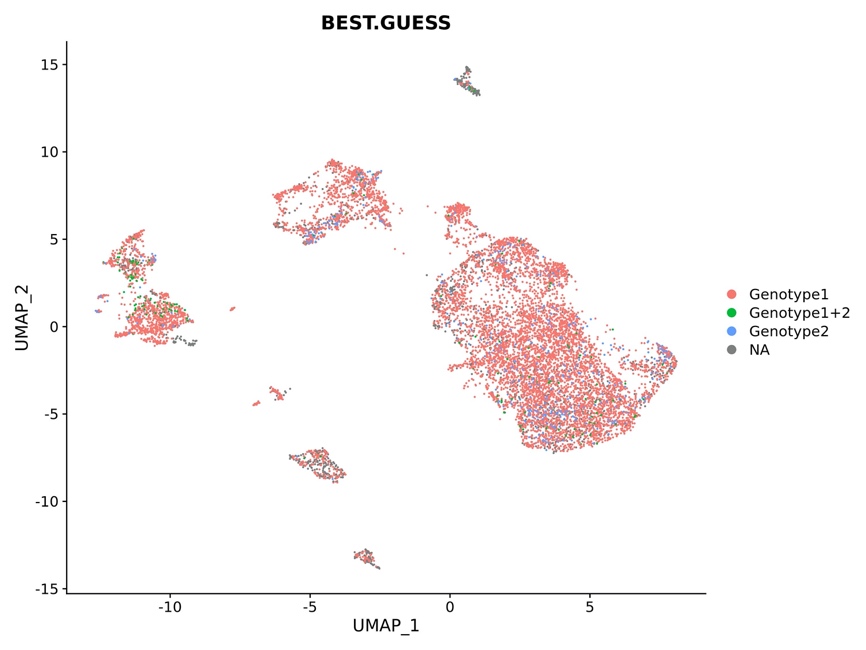

Supplementary Figure 8: Feature plots of conserved markers that are shared between the co-clustering trophoblast and cervical epithelial cells, plotted in the cervical (a) and placental (b) datasets.

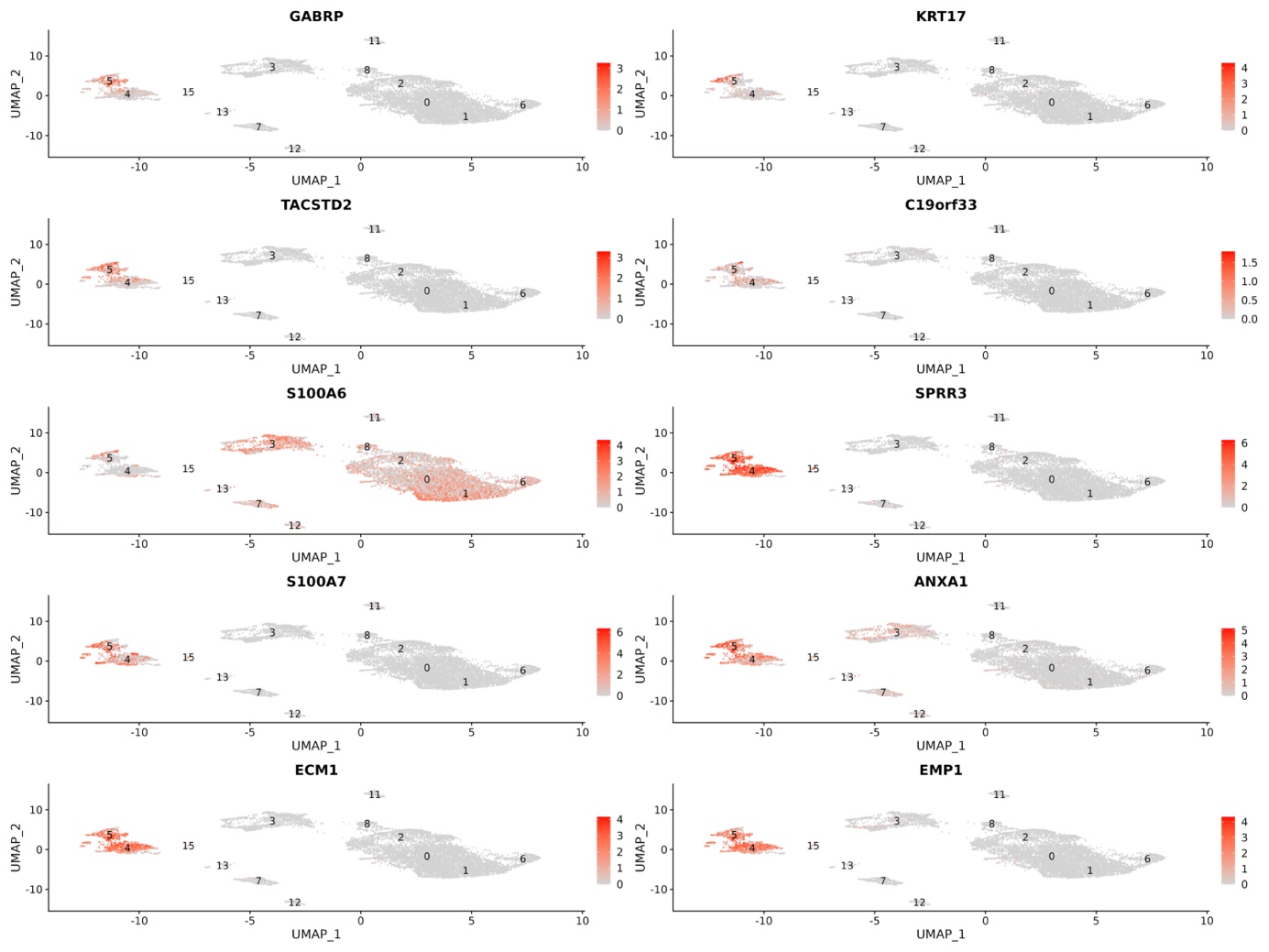

a

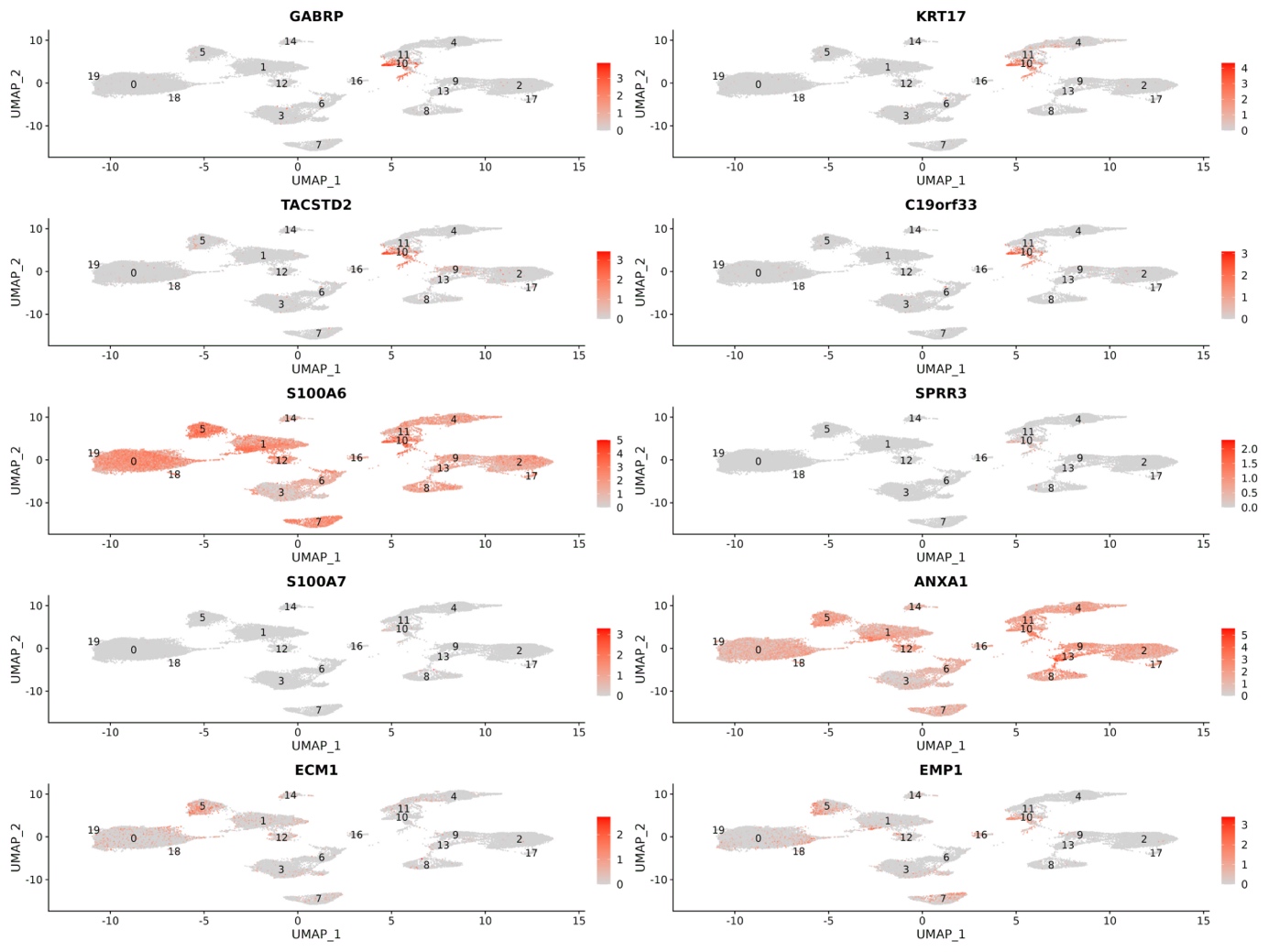

b

Supplementary Figure 9: Positive cells in CVS samples. (a) *HLA-G* positive cell, (b) *ADAM12* (green) and HLA-G (red), (c) *MCAM* (green) and *HLA-G* (red), (d) *PSG2* (green) and *HLA-G* (red).

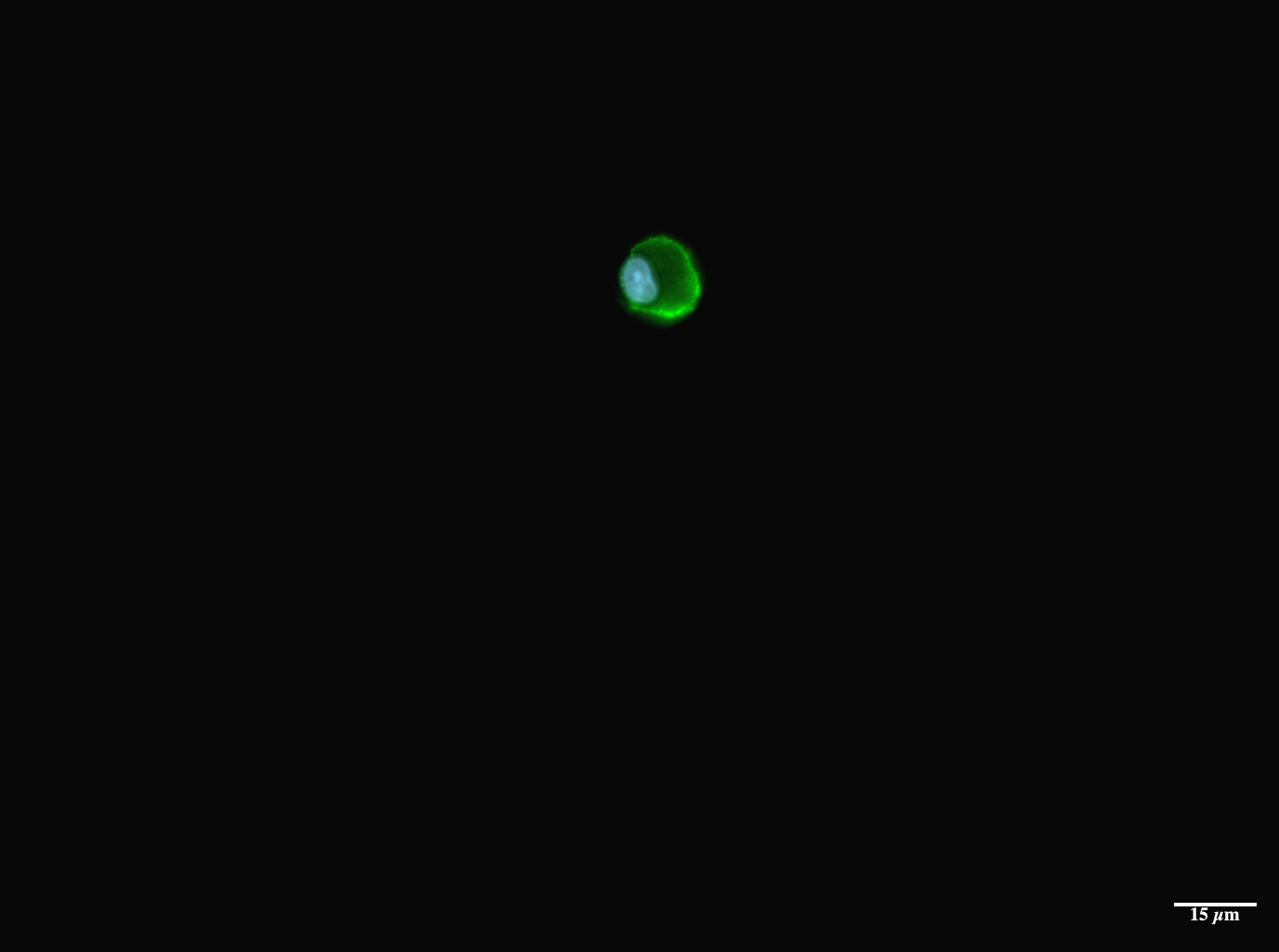

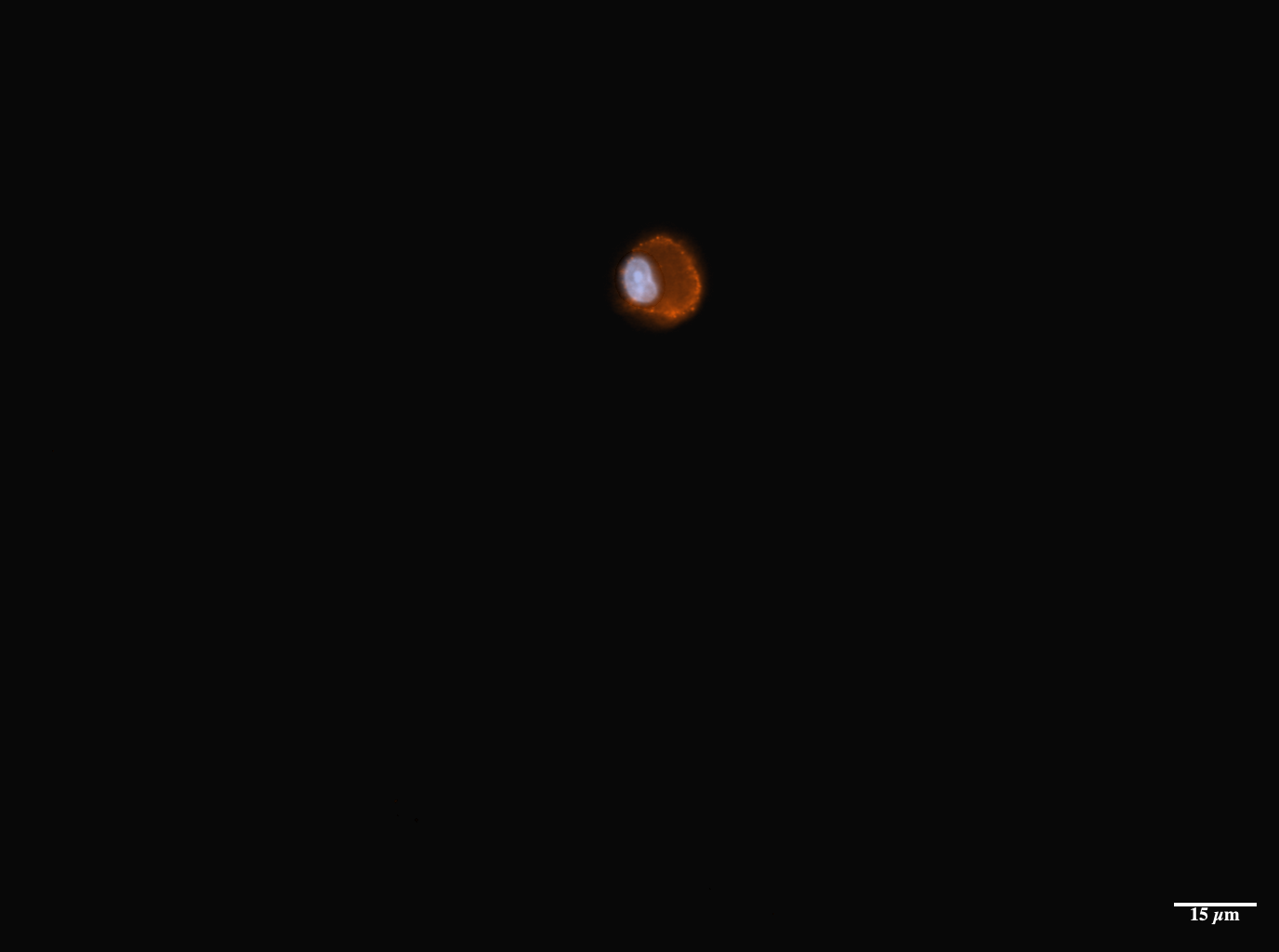

D

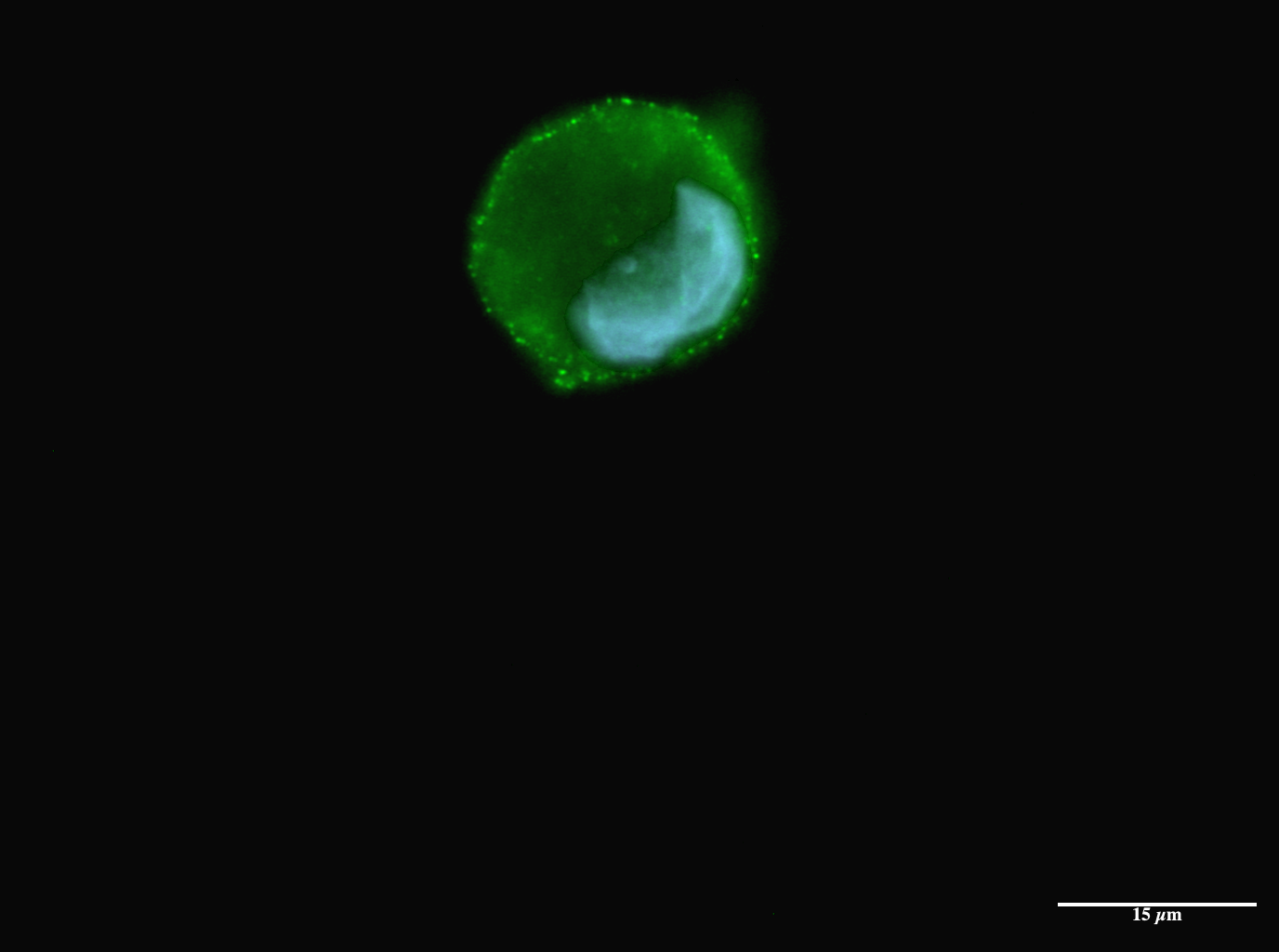

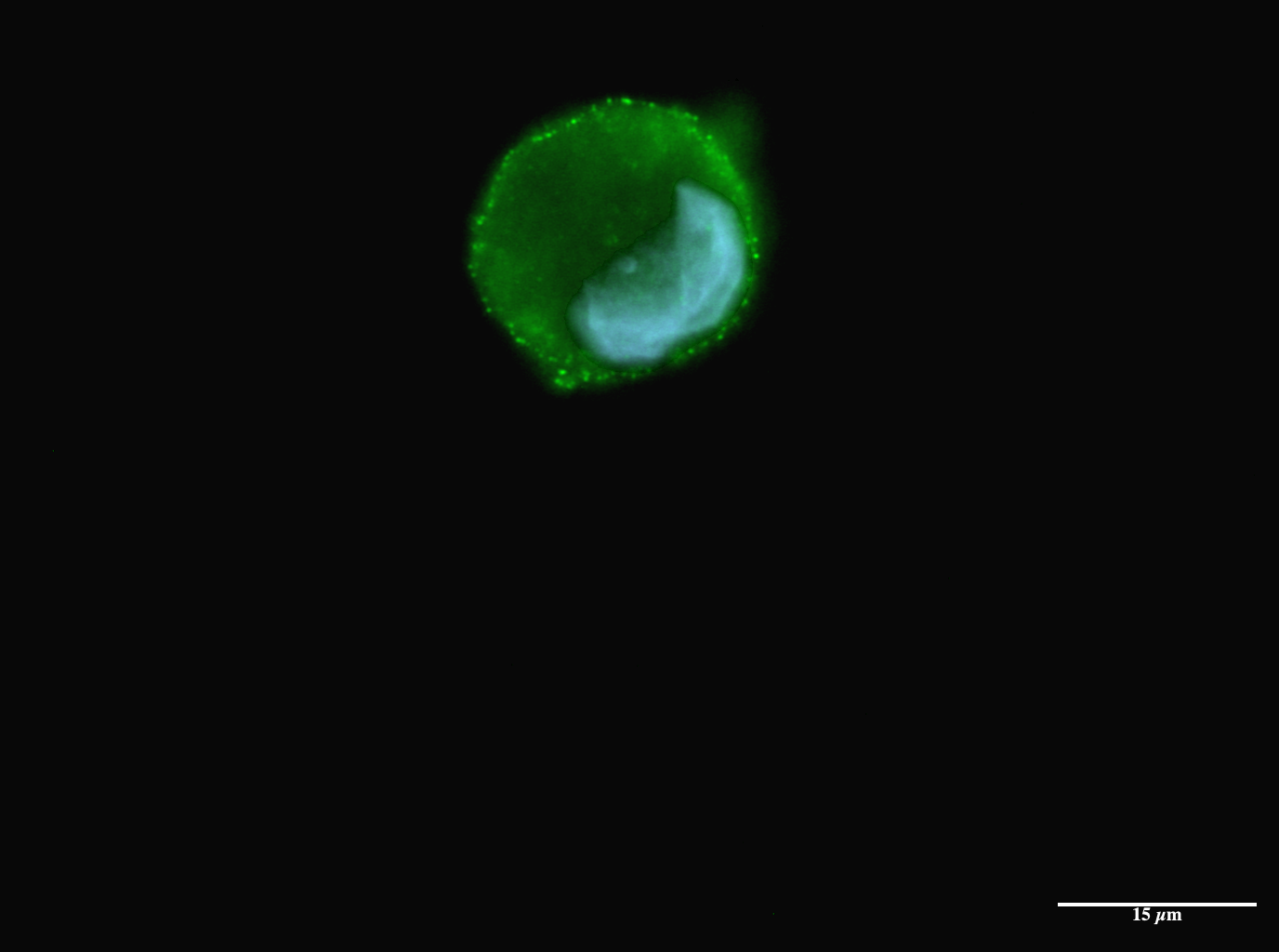

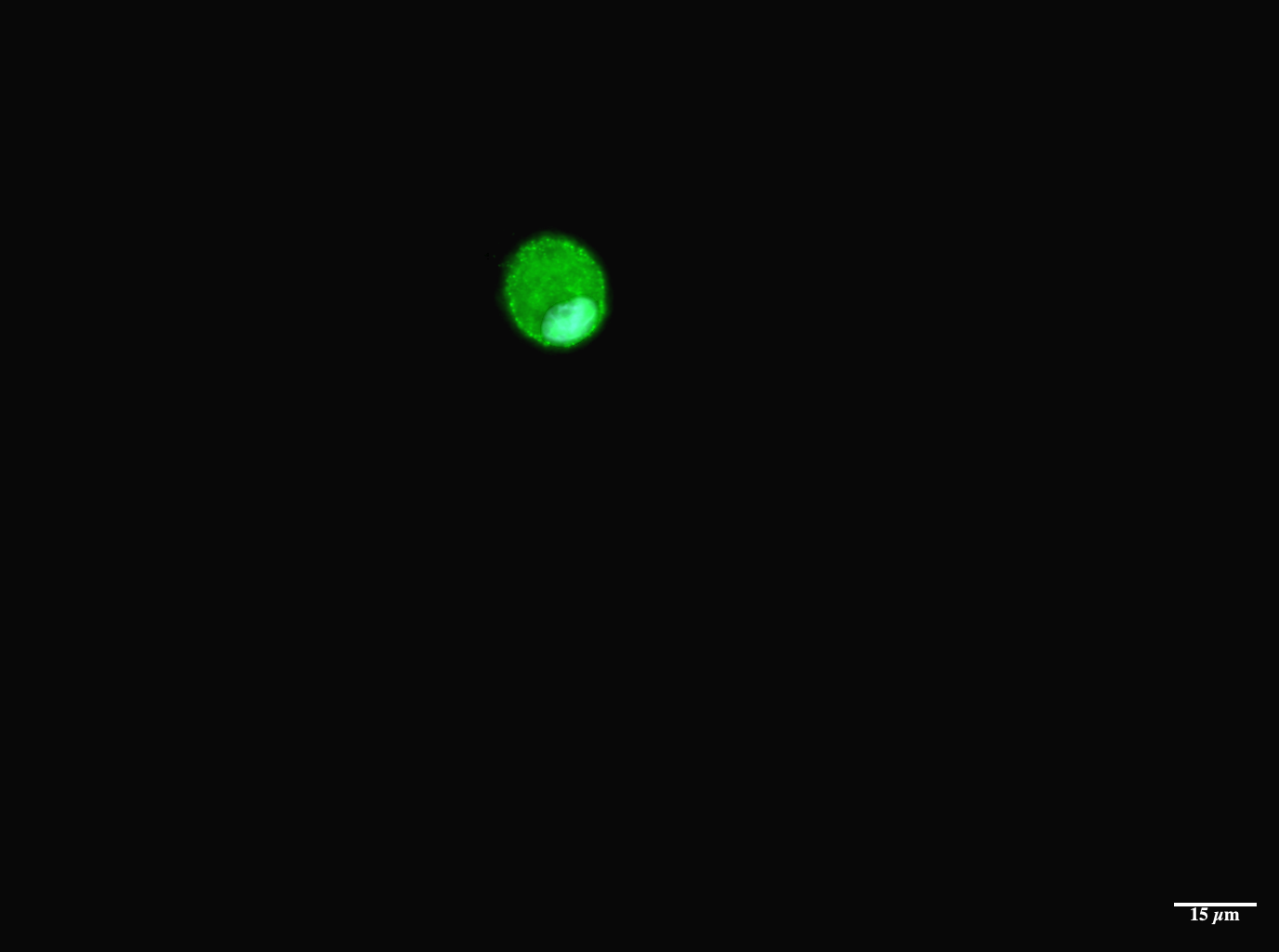

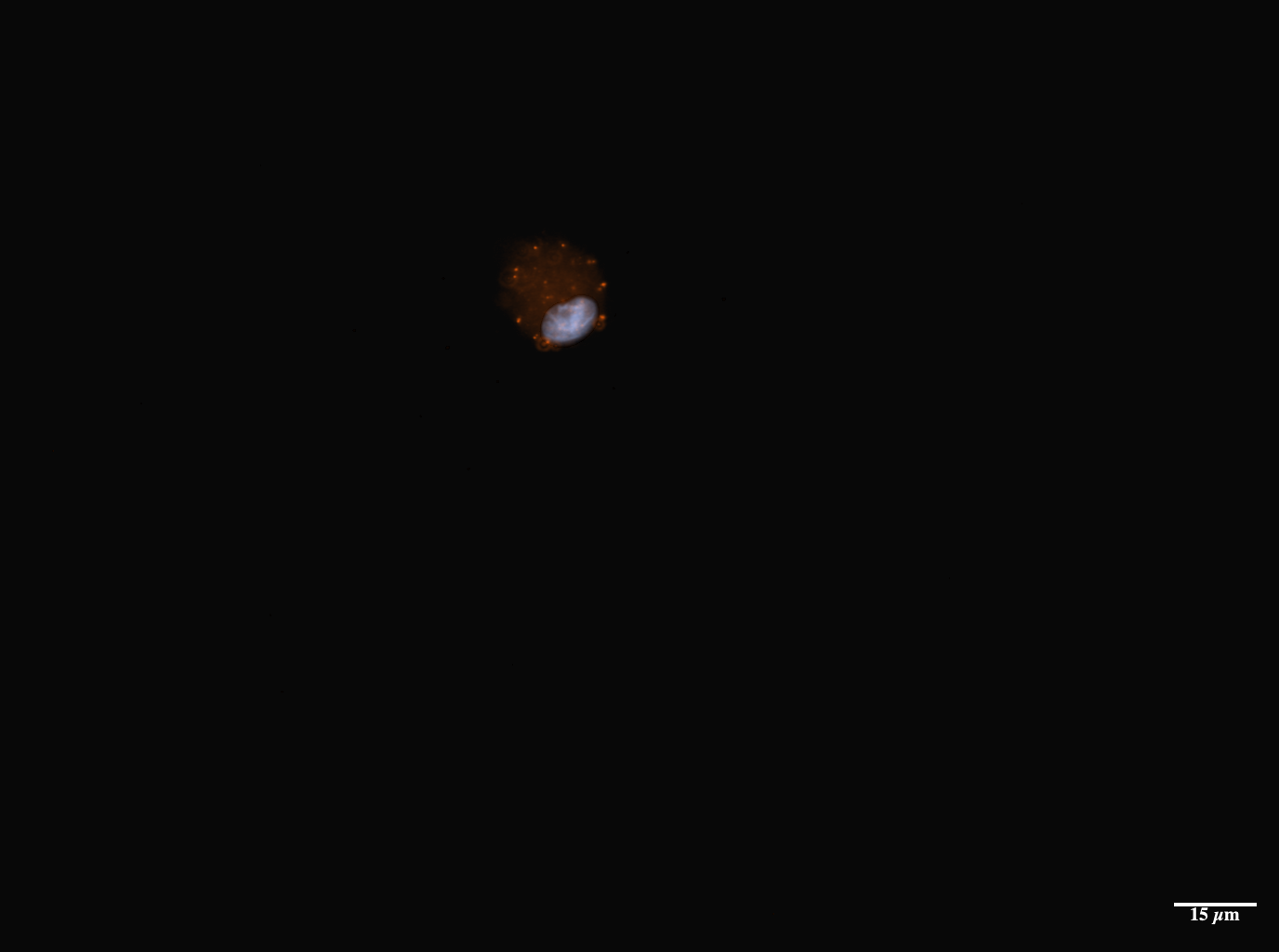

B

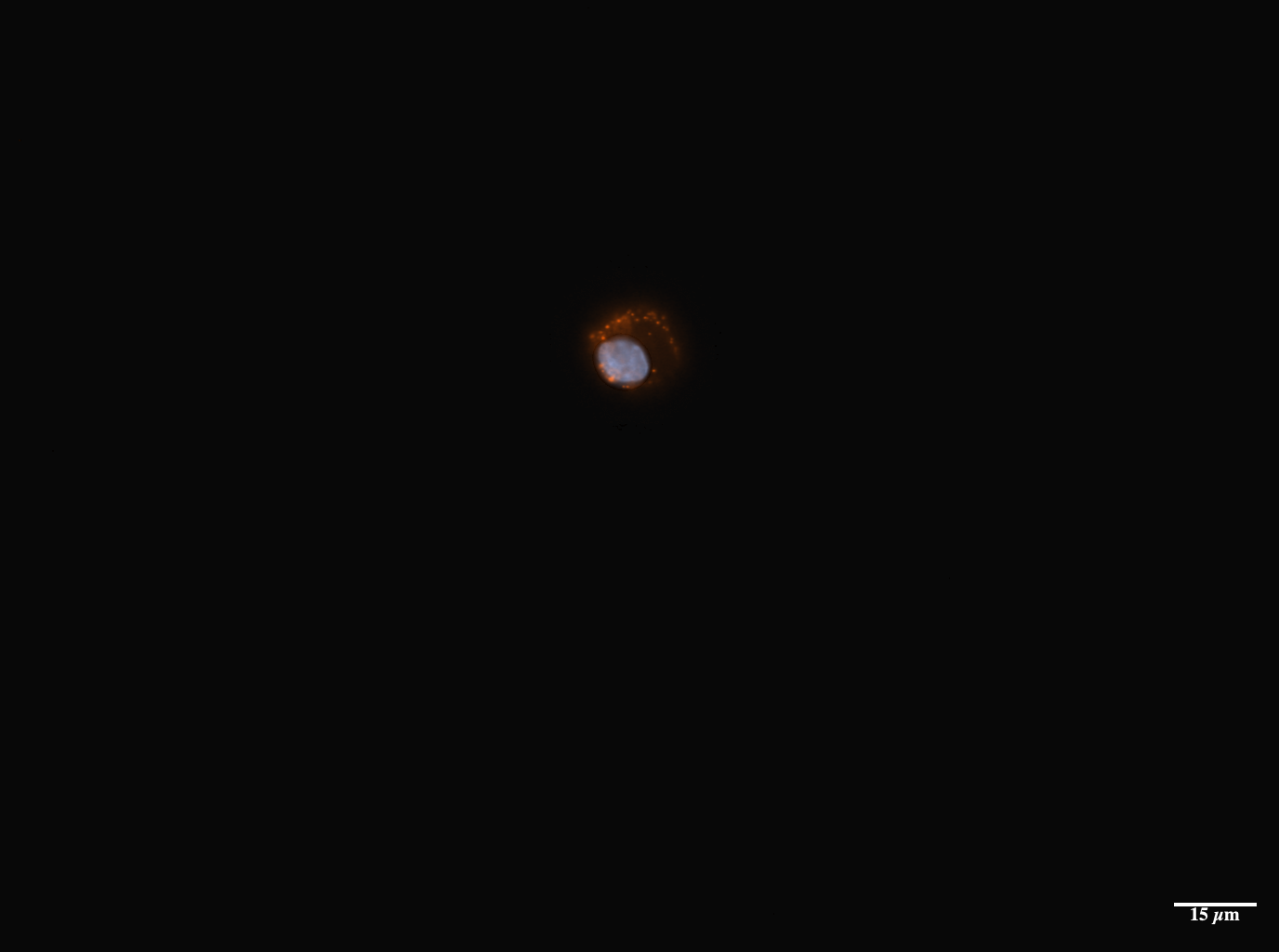

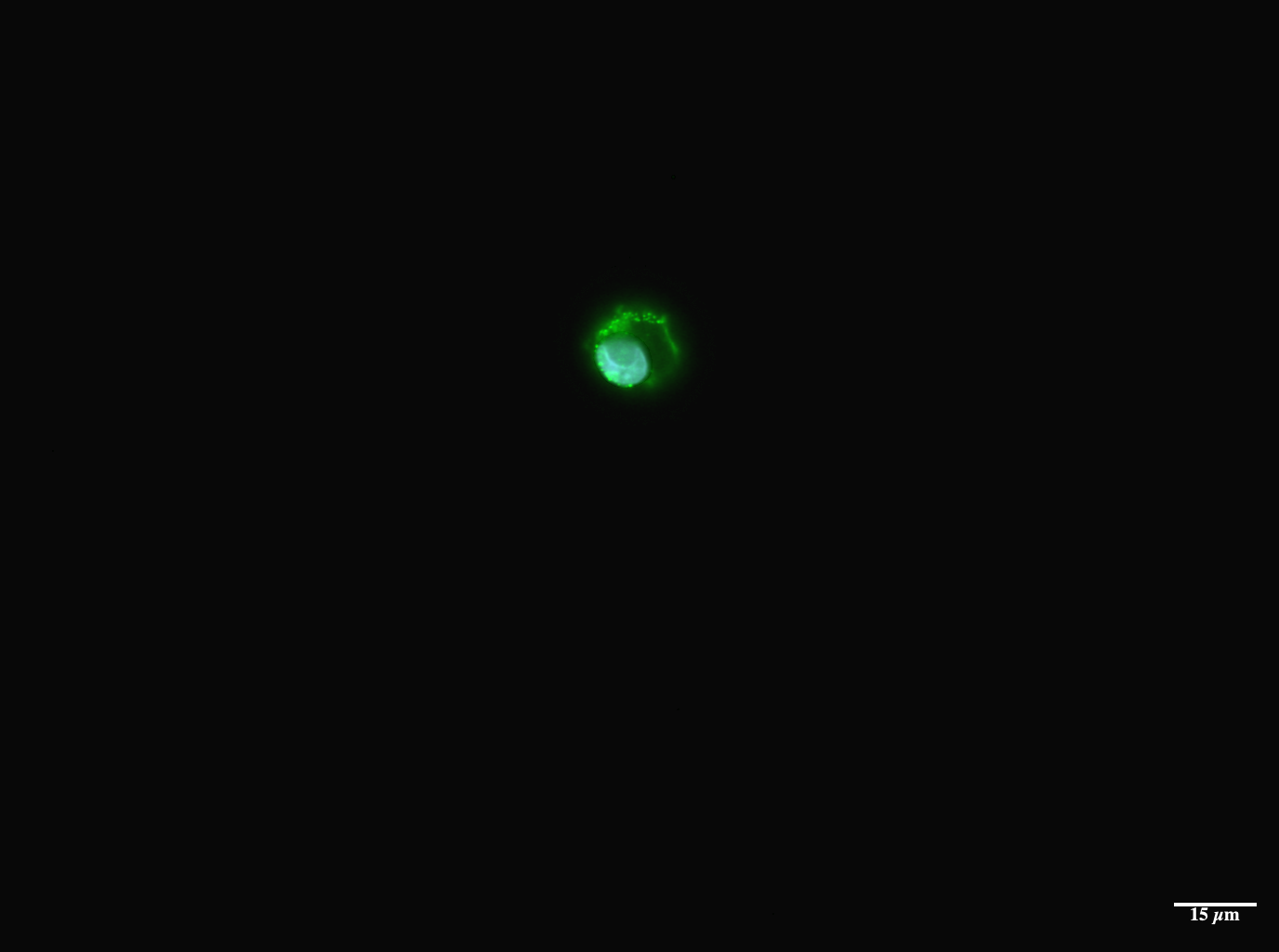

C

A

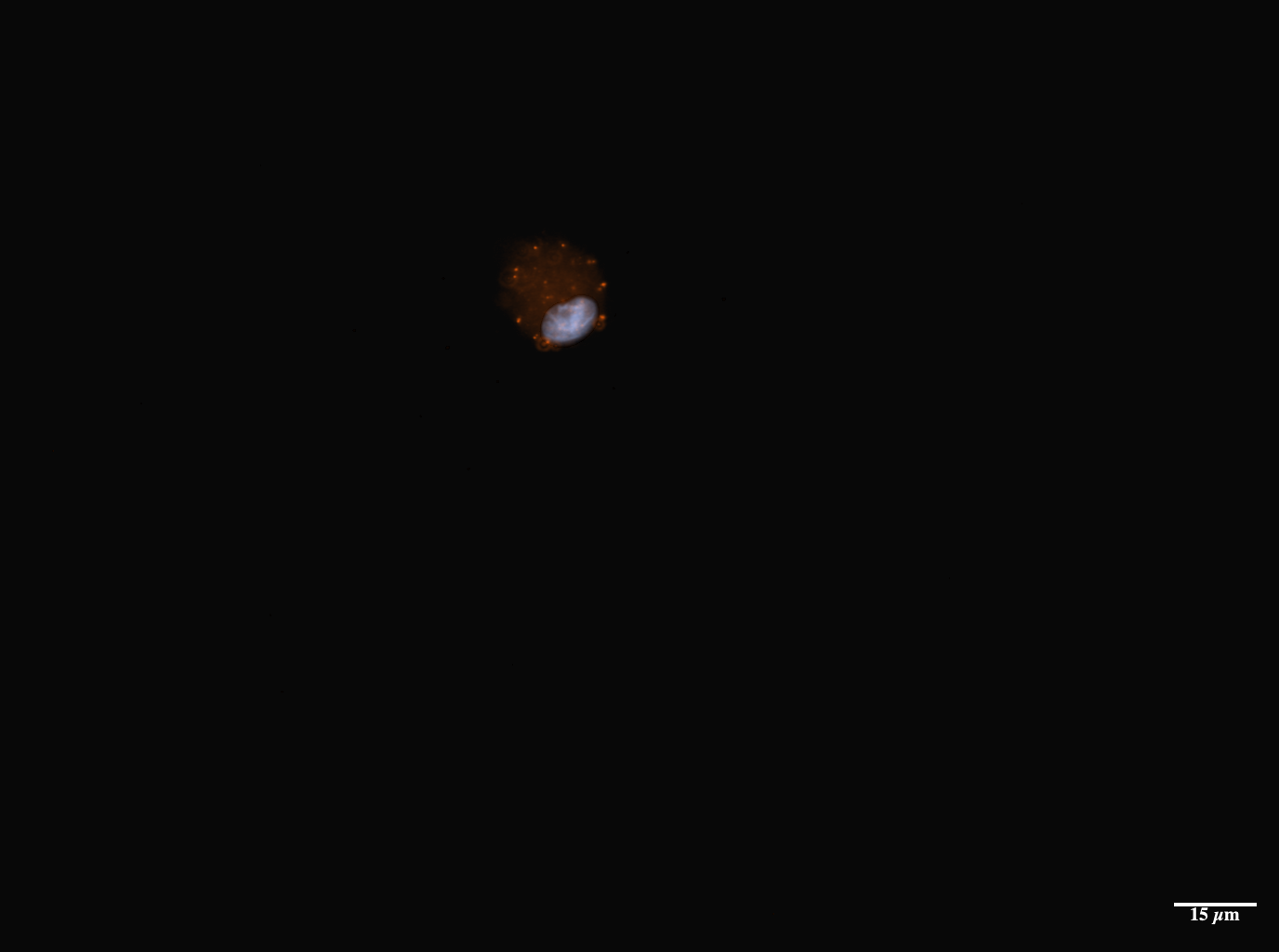

Supplementary Figure 10: Brightfield image of an exocervical sample on the LUNA-FL^TM^ Automated Cell Counter (Westburg). Large epithelial cells (asterisk) and smaller immune cells (arrow) are identified.

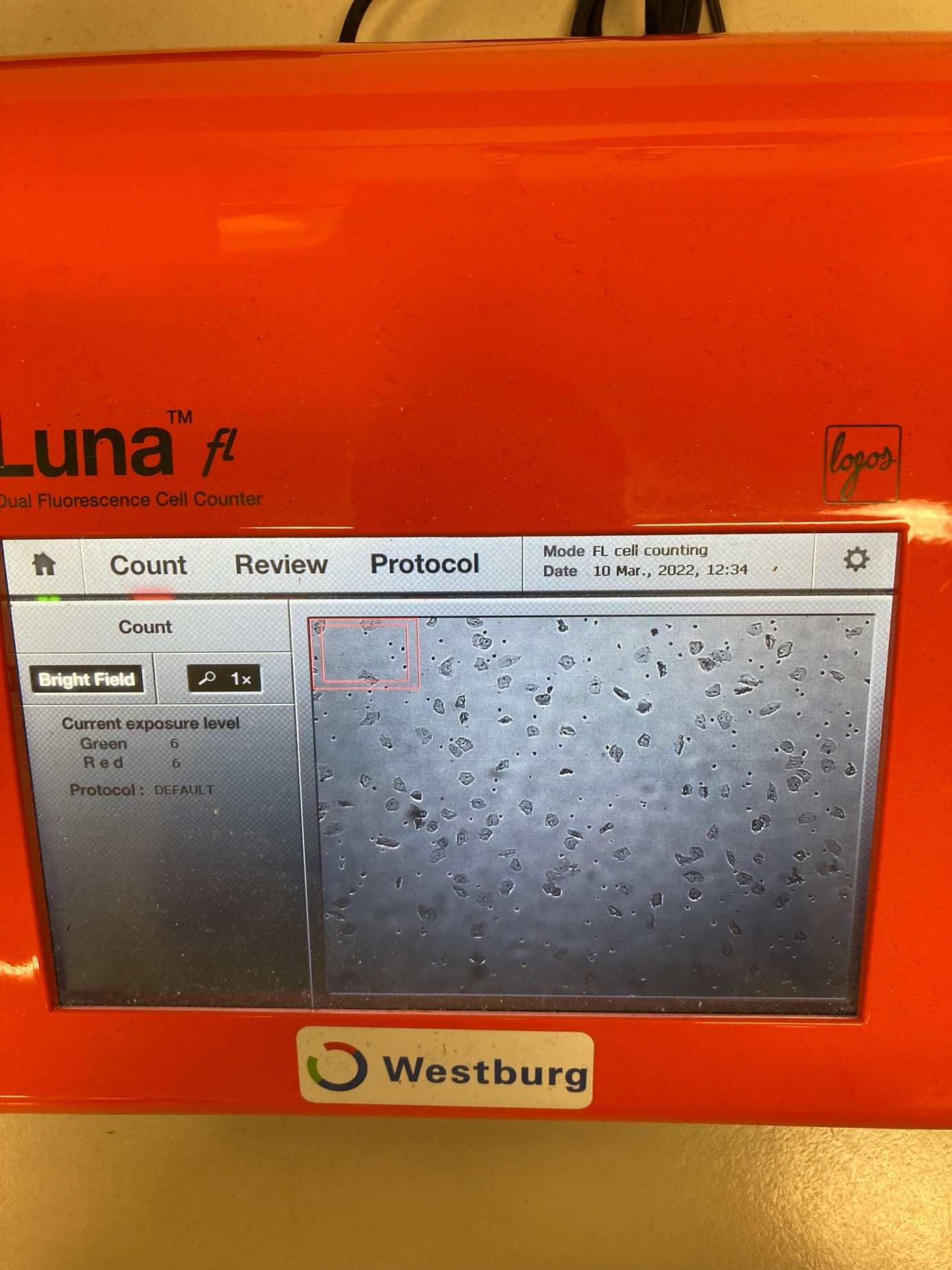

*

🡪

Supplementary Figure 11: Fluorescent signals in cervical epithelial cells. (a-b) background HLA-G staining when multiple epithelial cells form clumps. (c) cervical sample labeled in green with PSG2. Three epithelial cells are detected and each cell shows a variable intensity of fluorescent signal. Scale bar = 15 µm.

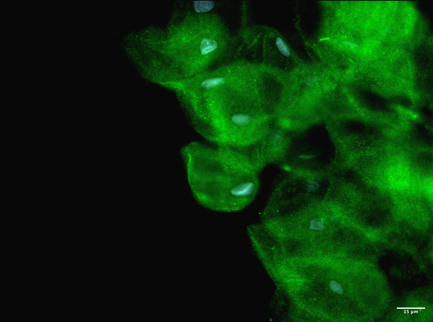

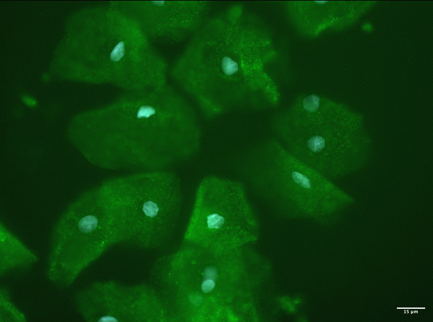

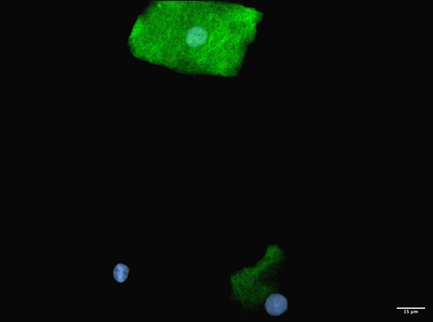

Supplementary Figure 12. Feature plot showing expression of general trophoblast marker *KRT7.* *KRT7* expression in (a) the epithelial cells of the exocervical dataset and (b) the trophoblast cells from the placental dataset. UMAP plots with the annotated cell types are depicted in (c).

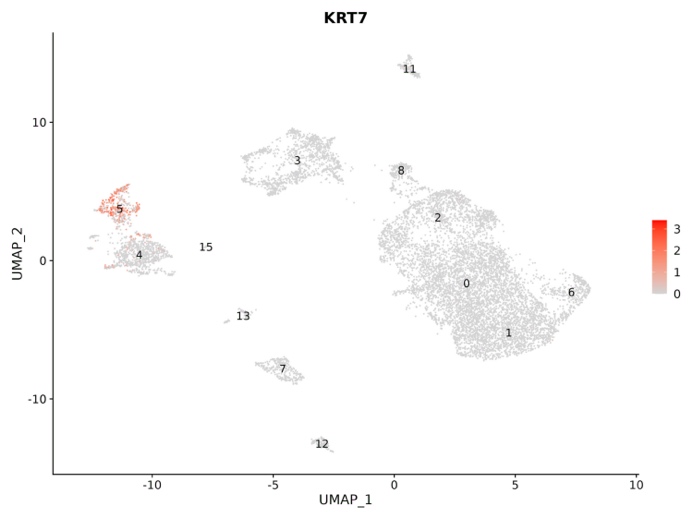

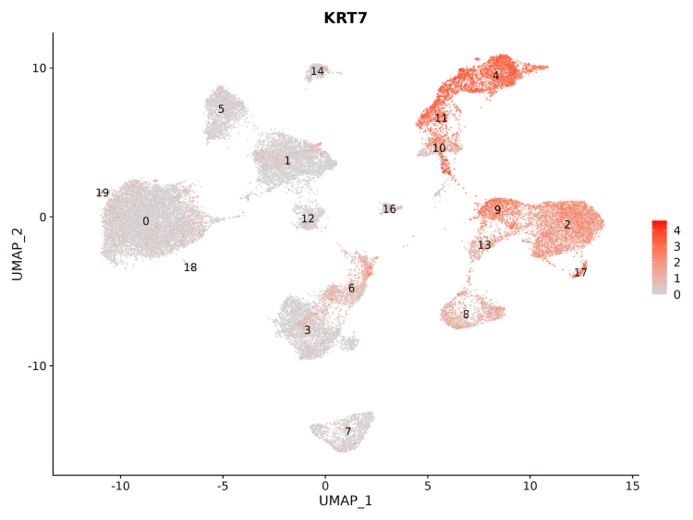

a

b

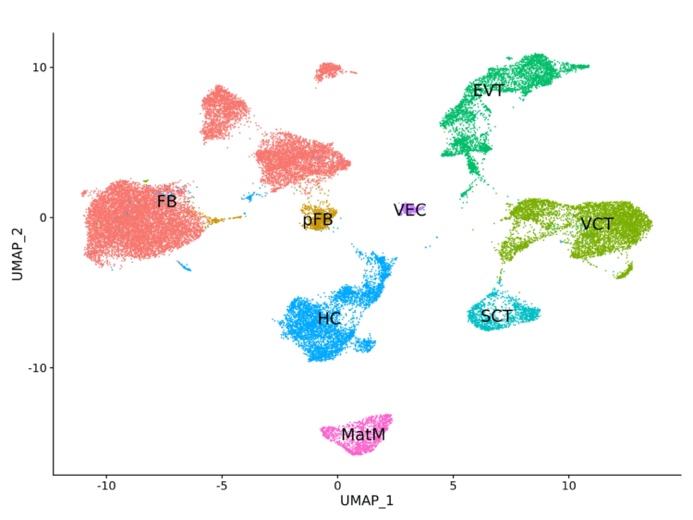

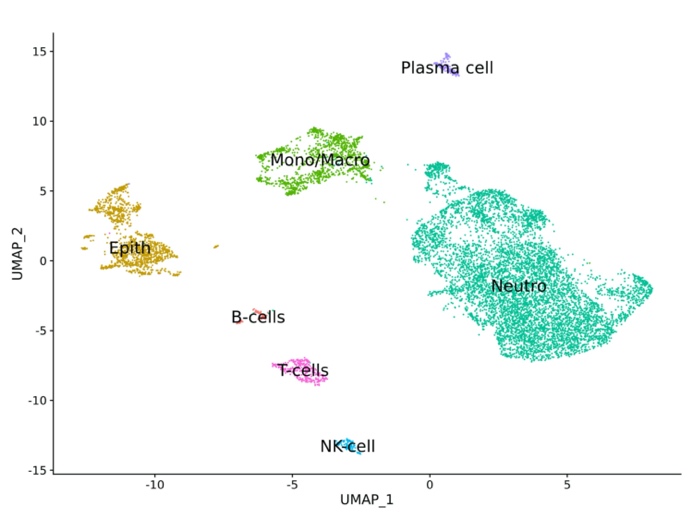

c

Supplementary Figure 13: Overview of laboratory procedures. Created with BioRender.com

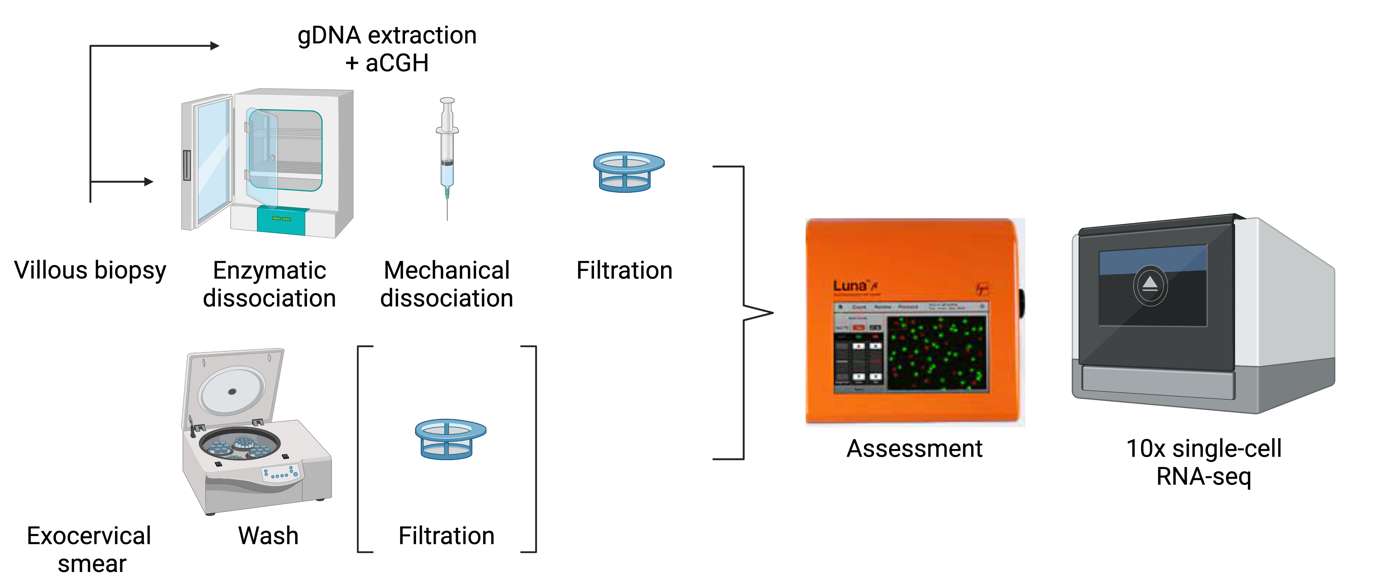

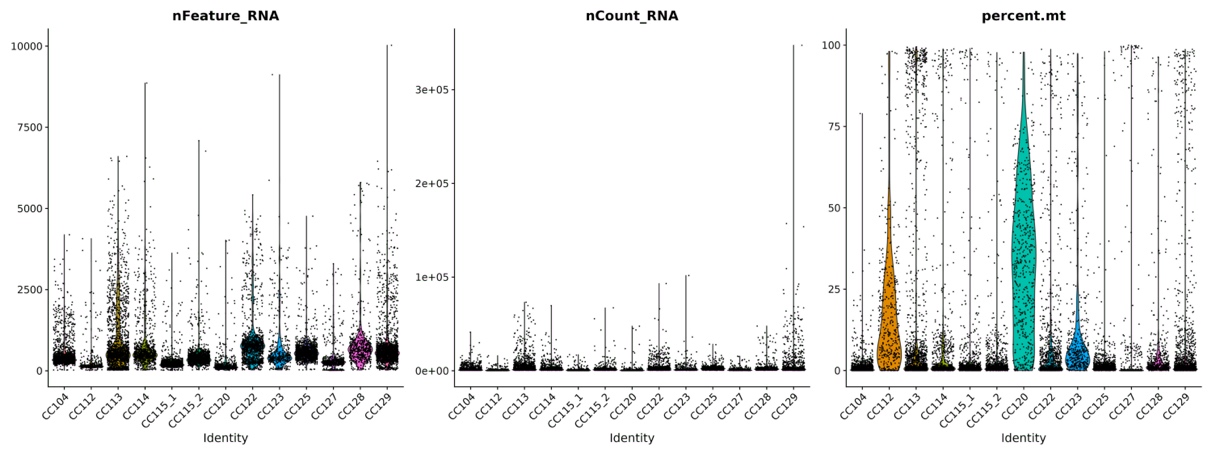
Supplementary Figure 14: Overview of QC metrics before (a) and after (b) filtering of the cervical dataset and placental dataset.

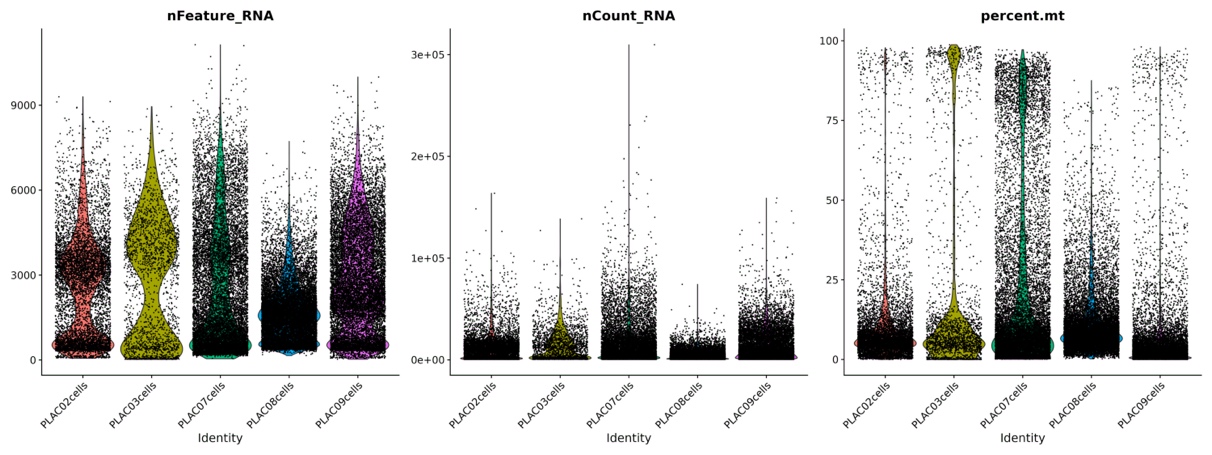

a

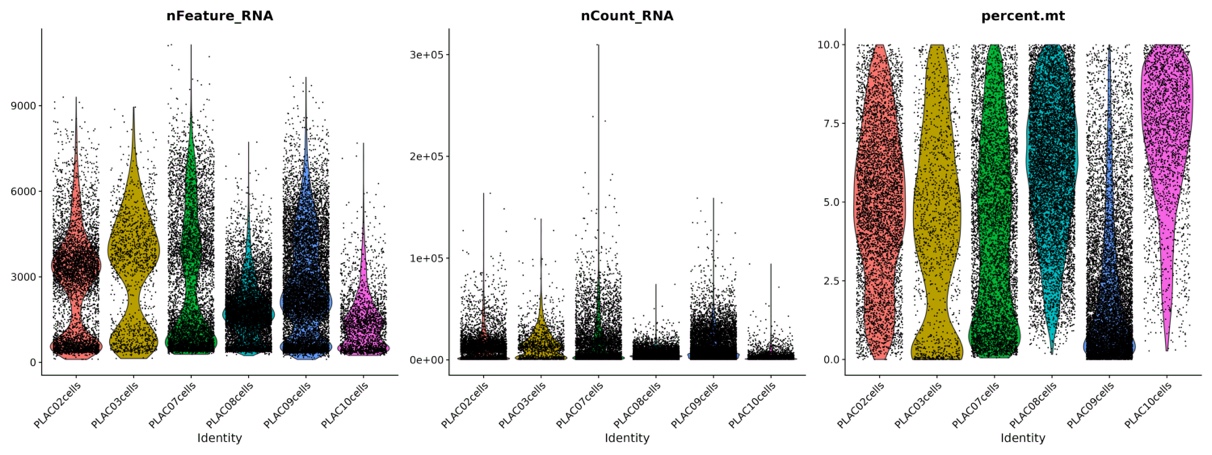

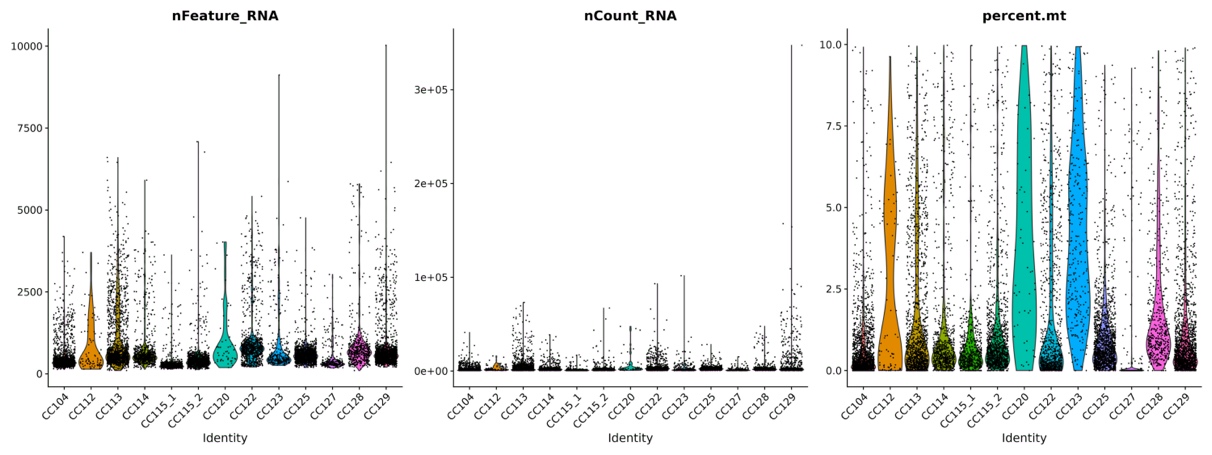

b

**Supplementary Tables**

Supplementary Table 1: Overview of the cervical samples included in the study.

Cell viabilities were determined by the LUNA-FL^TM^ Fluorescent Cell Counter (Westburg). Fetal sex was later defined by non-invasive prenatal screening. After CellRanger, filtered matrices were used as input for the analysis. Quality control and doublet removal were performed as described in the Methods section.

| Sample | Viability | Fetal sex | # cells (CellRanger output) | # cells after QC and doublet removal |
| --- | --- | --- | --- | --- |
| CC104 | 66.8% | Male | 1,905 | 1,813 |
| CC112 | 25.6% | Male | 419 | 85 |
| CC113 | 46.8% | Male | 1,765 | 1,354 |
| CC114 | 60.5% | Male | 893 | 734 |
| CC115 | 51.7% | Female | 1,275 and 1,273 | 616 and 1,175 |
| CC120 | 56.7% | Female | 541 | 53 |
| CC122 | 52.5% | Female | 1,189 | 965 |
| CC123 | 50.9% | Female | 443 | 291 |
| CC125 | 59.6% | Male | 1,578 | 1,517 |
| CC127 | 47.4% | Female | 550 | 258 |
| CC128 | 56.5% | Male | 554 | 483 |
| CC129 | 76.9% | Female | 1,928 | 1,629 |

Supplementary Table 2: The proportions of cell types for each exocervical sample.

Epith = epithelial cells, Neutro = neutrophils, Mono/Macro = monocytes and macrophages, NK-cells = Natural Killer cells.

| Sample | Epith | Neutro | Mono/Macro | T-cells | Plasma cells | NK-cells | B-cells |
| --- | --- | --- | --- | --- | --- | --- | --- |
| CC104 | 13,9% | 64,2% | 20,2% | 0,7% | 0,1% | 0,6% | 0,3% |
| CC112 | 13,3% | 51,8% | 27,7% | 3,6% | 0,0% | 2,4% | 1,2% |
| CC113 | 10,5% | 49,9% | 9,6% | 12,8% | 9,8% | 5,6% | 1,7% |
| CC114 | 6,5% | 79,1% | 9,6% | 1,0% | 0,3% | 0,7% | 2,8% |
| CC115_1 | 5,6% | 89,3% | 4,6% | 0,2% | 0,0% | 0,0% | 0,3% |
| CC115_2 | 16,1% | 79,3% | 3,5% | 0,5% | 0,2% | 0,1% | 0,3% |
| CC120 | 79,2% | 11,3% | 3,8% | 3,8% | 1,9% | 0,0% | 0,0% |
| CC122 | 15,1% | 68,4% | 11,7% | 3,1% | 0,3% | 0,6% | 0,6% |
| CC123 | 46,7% | 10,5% | 14,8% | 14,8% | 1,2% | 2,7% | 9,3% |
| CC125 | 0,9% | 94,3% | 4,3% | 0,1% | 0,1% | 0,1% | 0,1% |
| CC127 | 3,2% | 77,9% | 13,4% | 2,4% | 0,4% | 0,4% | 2,4% |
| CC128 | 13,2% | 78,0% | 6,4% | 0,6% | 0,4% | 1,3% | 0,0% |
| CC129 | 23,6% | 51,1% | 23,9% | 0,6% | 0,6% | 0,1% | 0,1% |

Supplementary Table 3: Placental samples included in the study.

The genotype is based on aCGH analysis of DNA extracted from a part of the placental biopsy. In parallel, aCGH was performed on DNA extracted from a fetal biopsy. The gestational age is calculated based on the date of the miscarriage diagnosis and the expected birth date. As input for the analysis, the filtered matrices of CellRanger were used. Quality control and doublet removal were performed as described in the methods sections.

| Sample | Placental genotype | Fetal genotype | Gestational age (weeks) | # cells (CellRanger output) | # cells after QC and doublet removal |
| --- | --- | --- | --- | --- | --- |
| PLAC02 | 47, XY +20 | Insufficient DNA | 9,7 | 7,284 | 5,218 |
| PLAC03 | 45, X | 45, X | 10,9 | 2,319 | 1,431 |
| PLAC07 | 47, XY +7 | 47, XY +7 | 11 | 13,500 | 6,436 |
| PLAC08 | 47, XY, +4 | XX (maternal contamination) | 9,9 | 15,317 | 8,888 |
| PLAC09 | 47, XX, +21 | 47, XX, +21 | 12,6 | 13,177 | 10,697 |
| PLAC10 | 46, XY | 46, XY | 10 | 5,361 | 2,271 |

Supplementary Table 4: Cell-specific marker genes of different placental cell types.

| Cell type | Clusters | Cell numbers | Marker genes |
| --- | --- | --- | --- |
| Fibroblasts | 0, 1, 5, 12, 14, 18, 19 | 16,636 | *COL1A1, DLK1, EGFL6* |
| Villous cytotrophoblasts (VCT) | 2, 9, 13, 17 | 5,742 | *PAGE4, PEG10, PHLDA2, XAGE3, VGLL1* |
| Extravillous trophoblasts (EVT) | 10, 11, 4 | 4,293 | *HLA-G, DIO2, LAIR2* |
| Syncytiotrophoblasts (SCT) | 8 | 1,245 | *CYP19A1* |
| Hofbauer cells | 3, 6 | 5,057 | *F13A1, CD14* |
| Maternal macrophages | 7 | 1,301 | *HLA-DRA, LYZ* |
| Vascular endothelial cells (VEC) | 16 | 291 | *PECAM1* |

Supplementary Table 5: Freemuxlet results of the final dataset of cervical cells.

The main genotype was labeled as Genotype (0,0), the alternative genotype as Genotype (1,1), and possible doublets as Genotype (1,0).

| Sample | Genotype (0,0) | Genotype (1,0) | Genotype (1,1) | % Genotype (1,1) |
| --- | --- | --- | --- | --- |
| CC104 | 1672 | 0 | 17 | 1.0% |
| CC112 | 81 | 0 | 2 | 2.4% |
| CC113 | 9 | 0 | 0 | 0% |
| CC114 | 683 | 0 | 7 | 1.0% |
| CC115.1 | 522 | 6 | 81 | 13.3% |
| CC115.2 | 1141 | 0 | 16 | 1.4% |
| CC120 | 53 | 0 | 0 | 0% |
| CC122 | 931 | 0 | 0 | 0% |
| CC123 | 256 | 1 | 1 | 0.4% |
| CC125 | 1472 | 3 | 9 | 0.6% |
| CC127 | 205 | 20 | 28 | 11.1% |
| CC128 | 400 | 27 | 30 | 6.6% |
| CC129 | 962 | 114 | 364 | 25.3% |

Supplementary Table 6: Mapping of the cells in the background cluster of the integrated dataset to the individual clustering results.

| Placental dataset | FB | pFB | HC | MatM | VCT | EVT | SCT | VEC |
| --- | --- | --- | --- | --- | --- | --- | --- | --- |
|  | 17 | 1 | 1694 | 10 | 388 | 510 | 17 | 15 |
| Cervical dataset | Neutro | Mono/Macro | NK-cells | B-cells | Plasma cells | T-cells | Epithelial cells |  |
|  | 120 | 11 | 0 | 28 | 137 | 1 | 11 |  |

Supplementary Table 7: Contributions of each sample to the clusters in the combined cervical and placental dataset.

|  | CC104 | CC112 | CC113 | CC114 | CC115_1 | CC115_2 | CC120 | CC122 | CC123 | CC125 | CC127 | CC128 | CC129 |
| --- | --- | --- | --- | --- | --- | --- | --- | --- | --- | --- | --- | --- | --- |
| 0 | 0 | 0 | 3 | 0 | 0 | 0 | 0 | 0 | 0 | 0 | 0 | 1 | 0 |
| 1 | 1123 | 43 | 654 | 538 | 551 | 928 | 5 | 619 | 14 | 1440 | 197 | 353 | 1133 |
| 2 | 0 | 0 | 13 | 0 | 0 | 0 | 0 | 0 | 1 | 0 | 0 | 0 | 0 |
| 3 | 0 | 0 | 0 | 0 | 0 | 0 | 0 | 0 | 0 | 0 | 0 | 0 | 0 |
| 4 | 45 | 0 | 24 | 9 | 0 | 0 | 0 | 7 | 0 | 4 | 1 | 2 | 5 |
| 5 | 20 | 1 | 143 | 26 | 0 | 6 | 2 | 24 | 23 | 3 | 2 | 12 | 16 |
| 6 | 352 | 24 | 139 | 86 | 26 | 27 | 2 | 118 | 68 | 36 | 41 | 27 | 83 |
| 7 | 242 | 11 | 104 | 43 | 36 | 197 | 42 | 150 | 120 | 16 | 8 | 63 | 360 |
| 8 | 0 | 0 | 2 | 0 | 0 | 0 | 0 | 0 | 0 | 0 | 0 | 0 | 0 |
| 9 | 0 | 0 | 0 | 0 | 0 | 0 | 0 | 0 | 0 | 0 | 0 | 0 | 0 |
| 10 | 0 | 0 | 0 | 2 | 0 | 3 | 0 | 1 | 1 | 0 | 0 | 0 | 0 |
| 11 | 0 | 0 | 0 | 0 | 0 | 0 | 0 | 0 | 0 | 0 | 0 | 0 | 0 |
| 12 | 0 | 0 | 0 | 0 | 0 | 0 | 0 | 0 | 0 | 0 | 0 | 0 | 0 |
| 13 | 0 | 0 | 1 | 0 | 0 | 0 | 0 | 0 | 0 | 0 | 0 | 0 | 0 |
| 14 | 24 | 6 | 263 | 20 | 2 | 6 | 2 | 34 | 60 | 1 | 9 | 9 | 10 |

| Cluster | PLAC02 | PLAC03 | PLAC07 | PLAC08 | PLAC09 | PLAC10cells |
| --- | --- | --- | --- | --- | --- | --- |
| 0 | 1515 | 443 | 1057 | 3996 | 2612 | 549 |
| 1 | 4 | 3 | 0 | 34 | 71 | 7 |
| 2 | 598 | 208 | 1381 | 609 | 1014 | 161 |
| 3 | 661 | 65 | 391 | 973 | 1639 | 61 |
| 4 | 348 | 142 | 22 | 306 | 1740 | 453 |
| 5 | 389 | 146 | 736 | 473 | 726 | 208 |
| 6 | 101 | 19 | 36 | 594 | 264 | 256 |
| 7 | 16 | 2 | 98 | 36 | 455 | 5 |
| 8 | 299 | 43 | 131 | 558 | 809 | 43 |
| 9 | 468 | 3 | 274 | 447 | 180 | 221 |
| 10 | 222 | 8 | 851 | 217 | 120 | 104 |
| 11 | 173 | 248 | 501 | 134 | 297 | 44 |
| 12 | 150 | 26 | 588 | 119 | 294 | 65 |
| 13 | 82 | 37 | 114 | 203 | 281 | 29 |
| 14 | 3 | 0 | 4 | 22 | 7 | 3 |
| 15 | 125 | 17 | 186 | 67 | 65 | 4 |
| 16 | 50 | 14 | 40 | 39 | 83 | 46 |
| 17 | 14 | 7 | 26 | 61 | 40 | 12 |

Supplementary Table 8: Potential membrane markers for EVT isolation from cervical samples.

*Specificity was assessed by plotting expression in the exocervical dataset.

| p_val | avg_log2FC | pct.1 | pct.2 | p_val_adj | gene | Unique for trophoblasts?* |
| --- | --- | --- | --- | --- | --- | --- |
| 0 | 3,37885567 | 0,871 | 0,017 | 0 | HLA-G | Yes |
| 0 | 2,97176929 | 0,997 | 0,426 | 0 | TPM1 | No |
| 0 | 2,86877264 | 0,761 | 0,013 | 0 | DIO2 | No |
| 0 | 2,85908222 | 0,849 | 0,078 | 0 | ADAM12 | Yes |
| 0 | 2,69283032 | 0,876 | 0,034 | 0 | PTPRF | No |
| 0 | 2,19583342 | 0,976 | 0,588 | 0 | ITGB1 | No |
| 0 | 1,98351201 | 0,829 | 0,108 | 0 | ITGA5 | No |
| 0 | 1,95598741 | 0,585 | 0,16 | 0 | TNFSF10 | No |
| 0 | 1,84958635 | 0,617 | 0,006 | 0 | MCAM | Yes |
| 0 | 1,72423798 | 0,993 | 0,932 | 0 | ITM2B | No |
| 0 | 1,69496722 | 0,792 | 0,162 | 0 | IFI6 | No |
| 0 | 1,50116672 | 0,749 | 0,013 | 0 | GJA5 | Yes |
| 0 | 1,49688576 | 0,526 | 0,031 | 0 | SFRP1 | Yes |
| 0 | 1,36561017 | 0,691 | 0,003 | 0 | CLDN19 | Yes |
| 0 | 1,35497281 | 0,395 | 0,009 | 0 | COL17A1 | Yes |
| 0 | 1,3168937 | 0,584 | 0,023 | 0 | IL2RB | No |
| 0 | 1,25312089 | 0,73 | 0,051 | 0 | NUCB2 | No |
| 0 | 1,20834771 | 0,68 | 0,081 | 0 | ENG | No |
| 1,456E-200 | 1,19547527 | 0,644 | 0,562 | 3,631E-196 | IL6ST | No |
| 0 | 1,1380117 | 0,994 | 0,858 | 0 | CD63 | No |
| 0 | 1,06544305 | 0,738 | 0,071 | 0 | CD151 | No |
| 0 | 1,03363819 | 0,551 | 0,044 | 0 | CAV1 | No |
| 0 | 1,00944684 | 0,787 | 0,169 | 0 | TMED10 | No |

Supplementary Table 9: Potential membrane markers for SCT isolation from cervical samples.

*Specificity was assessed by plotting expression in the exocervical dataset.

| p_val | avg_log2FC | pct.1 | pct.2 | p_val_adj | gene | Unique for trophoblasts?* |
| --- | --- | --- | --- | --- | --- | --- |
| 0 | 2,58366967 | 0,628 | 0,162 | 0 | IFI6 | No |
| 0 | 1,77771837 | 0,351 | 0,007 | 0 | PSG2 | Yes |
| 0 | 1,77219659 | 0,464 | 0,037 | 0 | EGFR | No |
| 0 | 1,61543397 | 0,643 | 0,168 | 0 | SLC40A1 | No |
| 0 | 1,59675423 | 0,611 | 0,008 | 0 | ERVW-1 | Yes |
| 0 | 1,47565441 | 0,382 | 0,078 | 0 | ADAM12 | Yes |
| 0 | 1,26827769 | 0,475 | 0,038 | 0 | FXYD3 | No |
| 0 | 1,17487106 | 0,346 | 0,025 | 0 | SMAGP | No |
| 0 | 1,14008867 | 0,386 | 0,003 | 0 | SLC22A11 | Yes |
| 0 | 1,12535916 | 0,369 | 0,051 | 0 | NUCB2 | No |
| 0 | 1,10570007 | 0,434 | 0,099 | 0 | APP | No |
| 0 | 1,0784292 | 0,501 | 0,021 | 0 | TUSC3 | No |
| 0 | 1,06775147 | 0,518 | 0,163 | 0 | SLC25A5 | No |
| 0 | 1,04859739 | 0,326 | 0,005 | 0 | BCAM | No |

Supplementary Table 10: Potential membrane markers for epithelial cell isolation from cervical samples.

*Specificity was assessed by plotting expression in the placental dataset.

| p_val | avg_log2FC | pct.1 | pct.2 | p_val_adj | gene | Negative in trophoblasts?* |
| --- | --- | --- | --- | --- | --- | --- |
| 0 | 4,27182503 | 0,625 | 0,001 | 0 | MAL | Yes |
| 0 | 3,67006069 | 0,612 | 0,034 | 0 | PSCA | No |
| 0 | 3,60995683 | 0,798 | 0,009 | 0 | EMP1 | No |
| 0 | 3,59628945 | 0,582 | 0 | 0 | RHCG | Yes |
| 0 | 3,05251861 | 0,574 | 0,001 | 0 | MUC21 | Yes |
| 0 | 2,41544239 | 0,487 | 0 | 0 | TMPRSS11B | Yes |
| 0 | 2,26833296 | 0,468 | 0,001 | 0 | TMPRSS11E | Yes |
| 0 | 2,24772984 | 0,402 | 0 | 0 | DUOX2 | Yes |
| 0 | 1,99992159 | 0,478 | 0,002 | 0 | MALL | Yes |
| 0 | 1,74379984 | 0,389 | 0,003 | 0 | TMPRSS2 | Yes |
| 0 | 1,72634395 | 0,403 | 0 | 0 | TMPRSS11D | Yes |
| 0 | 1,4915331 | 0,32 | 0 | 0 | CEACAM5 | Yes |
| 0 | 1,47670483 | 0,516 | 0,124 | 0 | GPRC5A | No |
| 0 | 1,46376177 | 0,394 | 0,013 | 0 | EPHA2 | No |
| 0 | 1,39948626 | 0,391 | 0,006 | 0 | MUC1 | No |
| 1,079E-126 | 1,29570542 | 0,245 | 0,064 | 2,69E-122 | CADM1 | No |
| 0 | 1,28349788 | 0,343 | 0 | 0 | CEACAM7 | Yes |
| 1,321E-276 | 1,23977223 | 0,261 | 0,015 | 3,295E-272 | MUC4 | No |
| 0 | 1,1768262 | 0,359 | 0,003 | 0 | SCNN1B | Yes |
| 1,695E-292 | 1,1460516 | 0,219 | 0 | 4,228E-288 | CLCA4 | Yes |
| 1,045E-209 | 1,12004384 | 0,284 | 0,042 | 2,606E-205 | F3 | No |
| 2,7264E-59 | 1,05154589 | 0,527 | 0,456 | 6,7996E-55 | B4GALT1 | No |

Supplementary Table 11: List of antibodies used for immunocytochemistry.

| Antibody | Final concentration | Host | Source |
| --- | --- | --- | --- |
| HLA-G (4H84) | 1/50 | Mouse | BD Biosciences **(557577)** |
| MCAM/CD146 | 1/250 | Rabbit | Abcam (ab75769) |
| ADAM-12 | 1/50 | Rabbit | ProteinTech  (141391-AP) |
| PSG2 | 1/100 | Rabbit | Abbexa (abx130865) |
| Goat anti-Mouse Alexa Fluor 488 | 1/600 | Goat | Life Technologies  (A-11001) |
| Goat anti-Mouse Alexa Fluor 568 | 1/600 | Goat | Life Technologies  (A-11004) |
| Goat anti-Rabbit Alexa Fluor 488 | 1/600 | Goat | Life Technologies  (A-11008) |
| Goat anti-Rabbit Alexa Fluor 568 | 1/600 | Goat | Life Technologies  (A-11011) |
